# High-Fidelity Tuning of Olfactory Mixture Distances in the Perceptual Space of Smell Through a Community Effort

**DOI:** 10.64898/2025.12.13.694160

**Authors:** Vahid Satarifard, Laura Sisson, Yikun Han, Pedro Ilídio, Matej Hladiš, Maxence Lalis, Xuebo Song, Wenjie Yin, Aharon Ravia, CiCi Xingyu Zheng, Gaia Andreoletti, Jake Albrecht, Robert Pellegrino, Zehua Wang, Stephen Yang, Robbe D’hondt, Achilleas Ghinis, Jasper de Boer, Felipe Kenji Nakano, Alireza Gharahighehi, DREAM Olfactory Mixtures Prediction Consortium, Benjamin Sanchez-Lengeling, Andreas Keller, Leslie B. Vosshall, Sébastien Fiorucci, Ambuj Tewari, Jérémie Topin, Celine Vens, Mårten Björkman, Danica Kragic, Noam Sobel, Nicholas A. Christakis, Joel D. Mainland, Pablo Meyer

## Abstract

A central goal in sensory science is to establish quantitative mappings between physical stimuli and perceptual responses. While such mappings are well characterized in vision and audition, they remain poorly defined in olfaction, limiting progress toward understanding the representations of smell. Predicting perceptual similarity between odor mixtures offers a promising route to formalize these relationships. To advance this effort, the DREAM (Dialogue for Reverse Engineering Assessment and Methods) Olfactory Mixtures Prediction Challenge assembled a curated, cross-study dataset describing the similarity of 507 mixture pairs and an unpublished test set of 46 mixture pairs. Teams competed to predict the perceptual similarity of mixture pairs, and then collaborated post-challenge to create an ensemble combining top-performing models that notably improves predictions over the existing state-of-the-art models. Moreover, ensemble model maintains high predictive accuracy in novel validation set. Our model provides a reproducible framework for neuroscientists, chemists, and engineers to compare odor mixtures and provides a foundation for future efforts towards better understanding the olfactory properties of mixtures.

## 1. Introduction

A central goal in sensory science is to discover how physical stimuli are represented in perceptual experience and to establish quantitative mappings between them. In vision, the trichromatic theory helps explain how combinations of three primary colors create our rich visual world. In audition, the superposition of sound waves at different frequencies produces complex tones and harmonies. These quantitative frameworks have enabled technologies from color television to digital audio, transforming how we capture, transmit, and reproduce sensory information. In olfaction, however, no comparable framework exists. We lack fundamental rules for predicting how molecular combinations produce odor perceptions, hindering both scientific understanding and technological applications in flavor and fragrance design.

However, recent advances in machine learning have made significant progress toward predicting the olfactory perception of single molecules from chemical structure. The first DREAM (Dialogue for Reverse Engineering Assessment and Methods) olfaction challenge [1] demonstrated that computational models could predict perceptual descriptors like “garlic,” “sweet,” and “floral”, achievements later expanded to broader descriptor sets [2]. The development of graph neural networks and larger datasets enabled construction of a Principal Odor Map (POM) [3] that maps molecular structures into perceptual space and predicts perceptual descriptors better than the median human panelist.

Most odors we encounter, however, are complex mixtures of dozens or hundreds of molecules. Although some models predict mixture perception from chemical structure [4–7], open competitions set clear benchmarks and often lead to improved predictions. With this motivation, we launched the second DREAM olfactory mixtures prediction challenge [8], inviting international teams to develop predictive models of perceptual similarity between pairs of molecular mixtures. For training, we curated a unified dataset from multiple psychophysical studies [4, 5, 9]. Over three months, 26 teams competed to minimize prediction error on a hidden test set of 46 mixture pairs, employing diverse machine-learning architectures and molecular feature representations.

The competition ended in a four-way tie among top performers. We combined predictions from the six highest-scoring teams into an ensemble model that outperformed all individual submissions and surpassed existing state-of-the-art methods, including the Snitz angle metric [4], POM-based approaches [3], semantic models [6], and aroma-chemical pair models [7]. The ensemble achieved a median root-mean-squared error of 0.08 and Pearson correlation of 0.57 on the test set. To validate these results, we generated an independent dataset of 50 mixture pairs and confirmed the model’s superior performance while establishing test-retest reliability benchmarks for human olfactory perception. Parsimonious models reveal the importance of perceptual features in accurate mixture predictions. By publishing all models in open-source format, we provide a quantitative, machine-readable framework for odor similarity that represents a crucial step toward understanding mixture perception and achieving the goal of capturing, transmitting, and reproducing olfactory information.

## 2 Results

### 2.1 The DREAM olfactory mixtures prediction challenge and dataset

For this competition, we standardized and compiled six datasets of odor-similarity measurements from three different studies [4, 5, 9] (see Methods and Table S1) into a unified dataset, comprising 168 unique mono-molecules, 731 unique mixtures, and 507 mixture pairs measurements. Mixture pair distances were mapped onto a continuous perceptual scale from 0 (indistinguishable) to 1 (maximally distinct), see Figure 1a left.

**Fig. 1.**
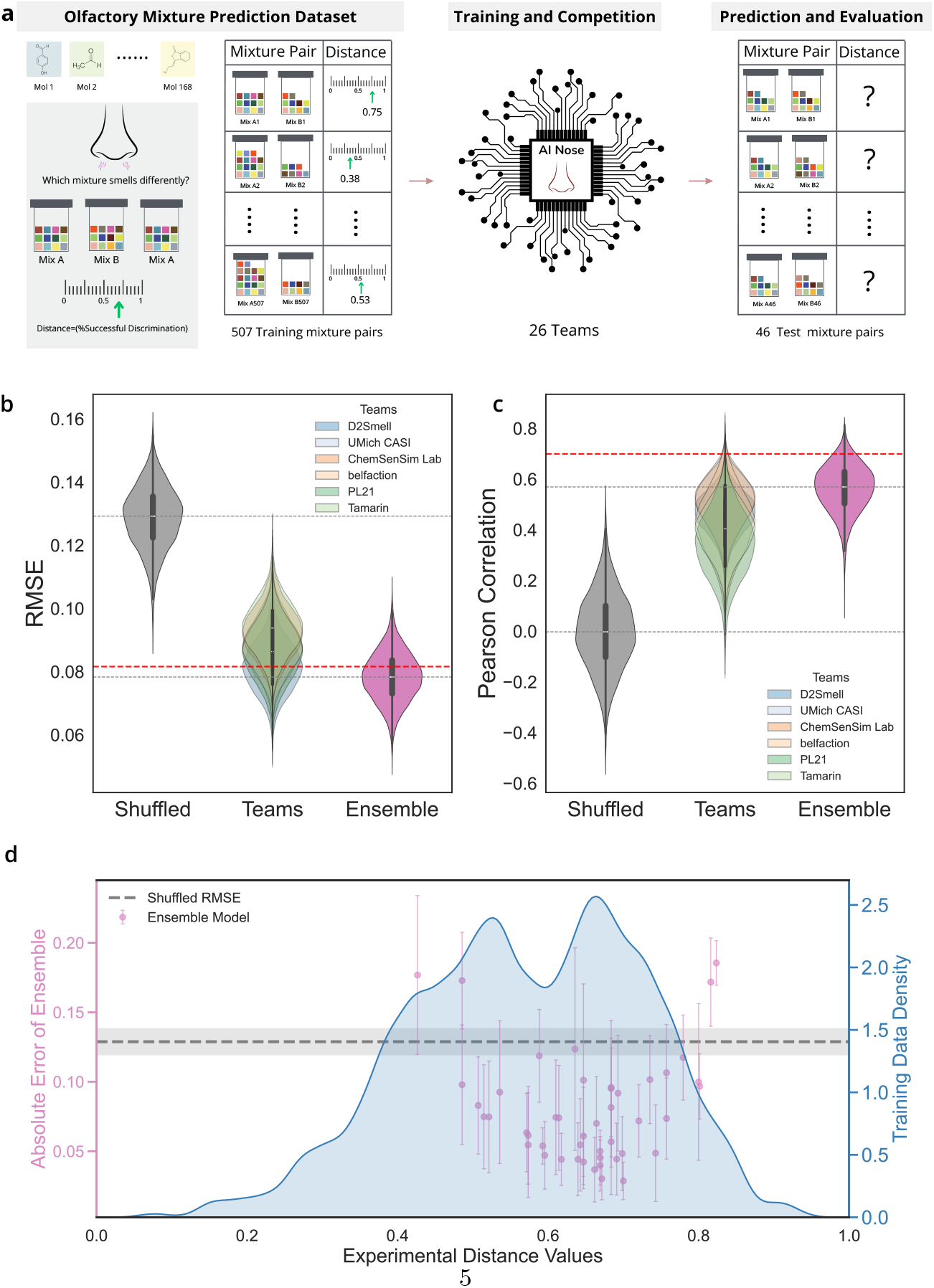
Overview of the challenge. **(a)** Schematic of the challenge workflow. *Left:* Participating teams received a curated training dataset of 507 olfactory mixture pairs with experimentally measured distances ranging from 0 (indistinguishable) to 1 (maximally distinct). *Middle:* Teams developed machine learning (ML) models to predict the perceptual distance of unseen mixture pairs, optimizing for minimal root mean square error (RMSE) and maximal Pearson correlation. The hidden test set comprised 46 mixture pairs. *Right:* 19 teams submitted final predictions for all 46 hidden test-set comparisons.**(b,c)** Distributions of RMSE **(b)** and Pearson correlation **(c)** for the six top-performing teams, the ensemble model, and a shuffled ground truth baseline, obtained via bootstrap sampling (n=10,000). Gray dashed lines indicate the median of the ensemble and shuffled distributions; red dashed lines denote within-subject split-half reliability (first half of the trials is used as test, and the second half as retest). **(d)** Absolute prediction error of the ensemble model (left y axis) as a function of experimental perceptual distance (x axis) overlaid with the training data density distribution (right y axis). The gray dashed line marks the shuffled baseline, and the gray band represents its standard deviation. Error bars show standard deviation across teams.

Teams competed for roughly three months,competition rules are presented in Methods, to develop machine-learning models to predict how close two molecular mixtures are in odor perceptual space, with the objective of minimizing the root mean squared error (RMSE) and maximizing Pearson correlation, respectively. In the competition phase of the challenge, teams were allowed to submit predictions for a Leaderboard holdout set of 46 pairs of mixtures, up to three times per week and receive feedback on the performance of their model. The ultimate goal of the challenge was to predict olfactory mixture distance for a hidden test-set of 46 mixture pairs (Figure 1a right). The test-set consisted of three subtypes: random, manual and angle. The random subtype contained mixture pairs generated without design constraints. The manual subtype included mixtures designed to be perceptually similar, and the angle subtype consisted of mixtures designed to be similar according to the Snitz angle distance metric [4]. Further methodological details are provided in Methods. The identities of all subtypes were concealed from participants throughout the competition.

DREAM organizers evaluated all submitted models using 10,000 bootstrap iterations, with RMSE and Pearson correlation as performance metrics, see Methods. The competition resulted in a four-way tie among the top-performing teams: *D2Smell, UMich CASI, ChemSenSim Lab*, and *belfaction*. We then built an ensemble model by averaging the predictions from these four teams with two additional high-performing models (*PL21*, and *Tamarin*, each achieving an RMSE below 0.1). A summary of these six models is presented in Table S2 with detailed descriptions of each model available in the Supplementary information. All other models were excluded either because of underperformance compared to random shuffled baseline, or they lacked open-source code for reproducibility, see Supplementary information and Figure S4. Distributions of RMSE and Pearson correlation for individual models and the ensemble are shown in Figure 1b,c. To estimate the performance ceiling imposed by intrinsic measurement noise, we calculated within-subject split-half reliability, where the first half of the trial measurements is used as test, and the second half is used as retest (red dashed lines in Figure 1b,c; see Figure S5). The ensemble model (pink distributions in Figure 1b,c) outperforms all individual models with a median RMSE of 0.08 and median Pearson correlation of 0.57 as indicated by gray dashed lines in Figure 1b,c. The ensemble model performed best on manual and angle subtypes, with Pearson correlation of 0.38 for both, and RMSE of 0.07 and 0.06, respectively. For the random subtype, it achieved an RMSE of 0.10 and slightly higher Pearson correlation of 0.45 (see Figure S7 for subtype distributions).

Analysis of the residuals (the difference between model predictions and ground truth) for each test-set measurement shows that models generally tend to underestimate perceptual similarity (Figure S7). All teams except *PL21* exhibit a net negative residual sum. The random subtype also showed larger residuals than the manual and angle subtypes. Among all models, *PL21* displayed a distinct performance profile, performing relatively well on the random subtype but poorly on the manual and angle subtypes (see Figure S7).

Model performance also correlated with the density of training data (Figure 1e). Prediction error is lowest for mixtures that were rated in the midrange (approximately 0.5 and 0.8), where data are most abundant. In contrast, errors increase for distances below 0.5 or above 0.8, likely reflecting the scarcity of training examples in these regions.

### 2.2 The ensemble model outperforms the state of the art

To evaluate the performance of our ensemble model, we compared it with four state-of-the-art (SOTA) models: the Snitz angle model [4, 5], the open-POM model (an open source implementation of POM [3, 10]), a semantic model [6], and an aroma chemical pair model [7]. Computational details are provided in Methods and Figure S8.

To estimate the distributions of RMSE and Pearson correlation we performed bootstrap sampling (n=10,000)(see Methods). To establish a chance-level baseline, we shuffled the experimental distance values. Figure 2a,b shows the resulting distributions for the shuffled ensemble, individual SOTA models (linear fits) and the ensemble model. All SOTA models outperformed the random shuffle baseline (see Figure 2a) and all achieved Pearson correlations above chance level, with the ensemble model performing best (Figure 2b). Across test-set subtypes, the largest improvements occurred for the manual and random subtypes: RMSE decreased by 0.039 *±* 0.004 and 0.047 *±* 0.005 while Pearson correlation increased by 0.29 *±* 0.07 and 0.26 *±* 0.10, respectively (Figure S9). When averaged across all subtypes, the ensemble model reduced RMSE by 0.035 *±* 0.004 and increased Pearson correlation by 0.22 *±* 0.05. Overall, the ensemble model consistently outperformed all SOTA models and combining SOTA predictions with the ensemble models decreased the performance (Figure S9).

**Fig. 2.**
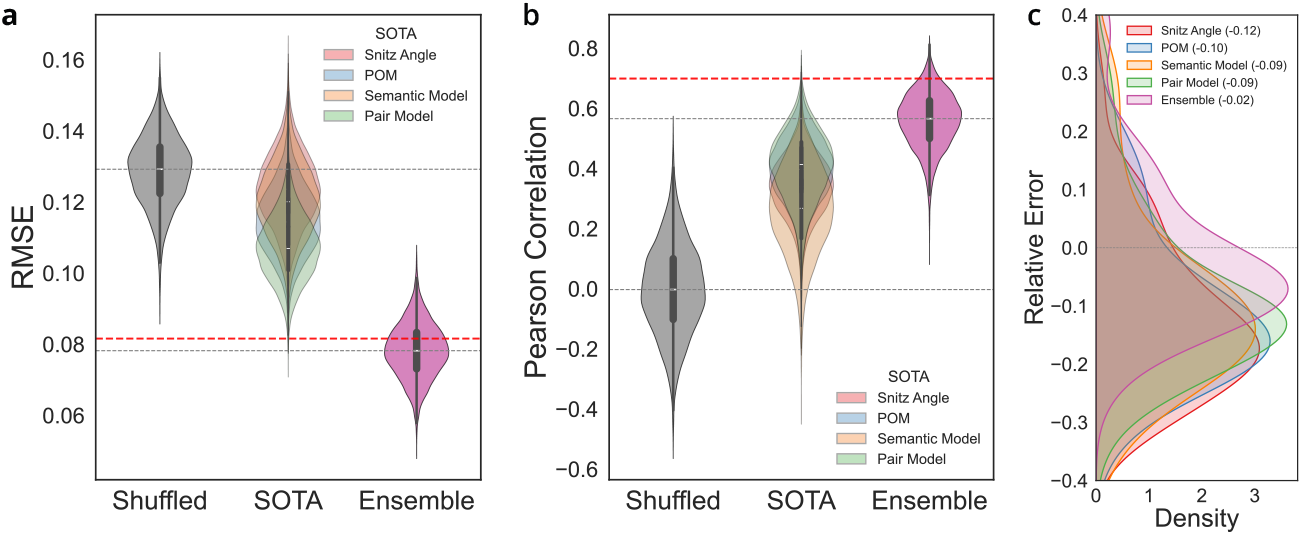
The ensemble model outperforms existing state-of-the-art approaches. **(a,b)** Distributions of RMSE **(a)** and Pearson correlation (y axis)**(b)** for the ensemble model (pink), individual state-of-the-art (SOTA) models and a shuffled ground-truth baseline (gray) obtained via bootstrap sampling (n=10,000). Gray dashed lines indicate median values for the ensemble model and shuffled baseline, and the red dashed line marks the across-subject split-half reliability (first half as test, second half as retest). **(c)** Kernel density estimates of relative-error distributions for each SOTA model and the ensemble model on the test set. The dashed line at 0 indicates no bias; legend values report the mean values of relative error for each model.

We next analyzed relative error, defined as 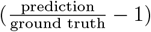 for each SOTA model and the ensemble model ( see Figure S10). Across all models and test-set subtypes, relative error showed a negative correlation with experimental distance values. This pattern indicates that all models tend to bias predictions towards intermediate values, overestimating lower perceptual distances and underestimating higher ones. The increase in relative error at smaller distances is expected, since dividing by small ground truth values amplifies relative differences. Error growth at both extremes also reflects limited training data in these regions (Figure 1d). As shown in Figure 2c, the ensemble model’s relative error distribution is shifted toward lower errors compared with the SOTA models, with a mean value of -0.02 *±* 0.13.

### 2.3 Optimal features emerge from feature importance analysis

To quantify feature importance across six different models we conducted a SHAP (SHapley Additive exPlanations) value analysis [11, 12] (Figure 3). Each model included over 100 individual features, so we grouped them into feature sets for interpretability (x-axis labels in Figure 3a). For each set, feature importance was defined as the mean absolute SHAP value across test-set predictions; error bars represent the standard deviation of these means (x-axis in Figure 3a). Detailed definitions of the feature sets used in each model are provided in Supplementary information.

**Fig. 3.**
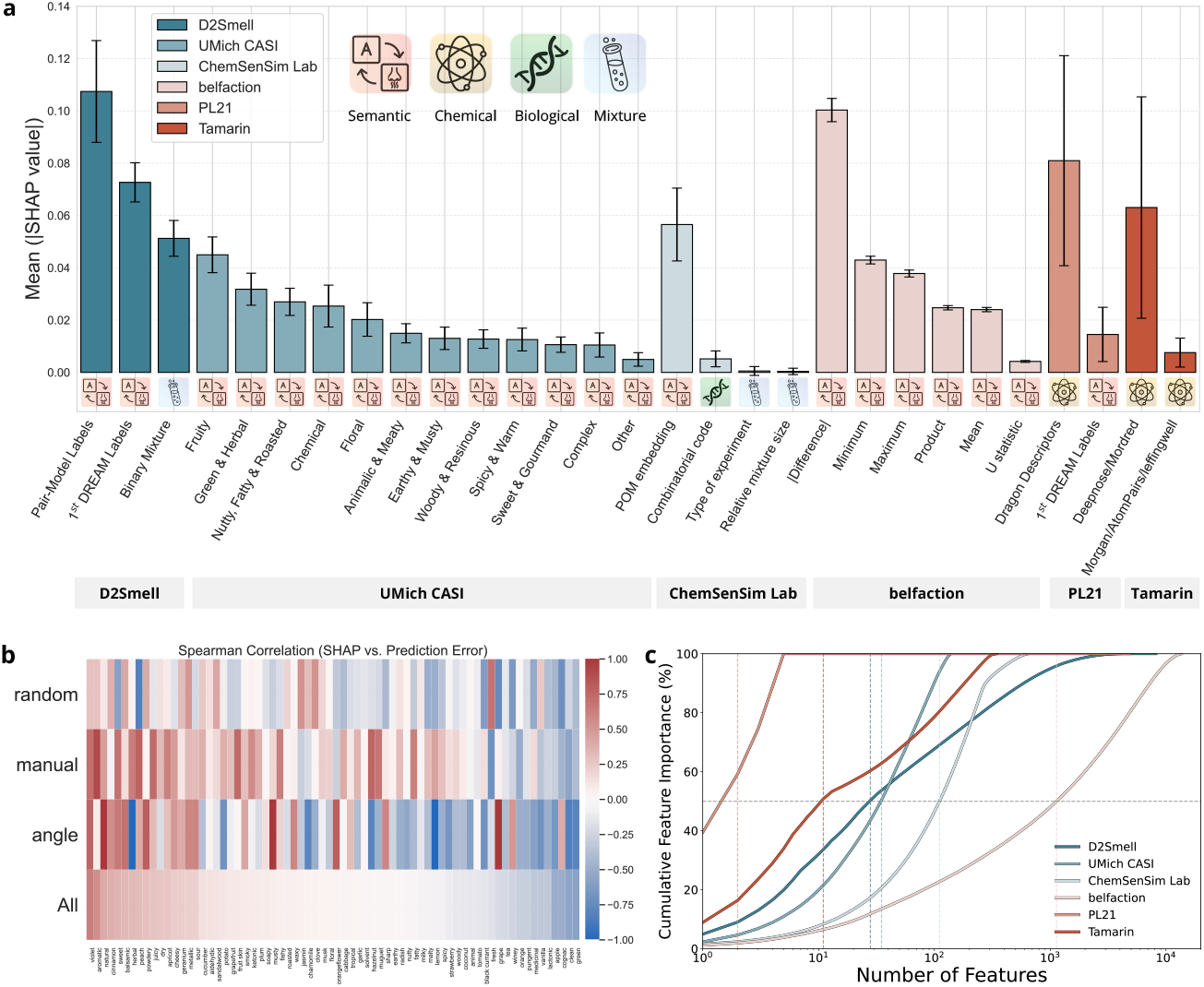
Feature importance analysis of top-performing models. **(a)** Mean absolute SHAP (SHapley Additive exPlanations) values for grouped feature classes across six teams. Larger values indicate greater contribution to model predictions. Error bars represent the standard deviation. Different class of features (semantic, chemical, biological, and mixture), are differentiated with different symbols in the figure inset.**(b)** Heatmap of Spearman’s rank correlations between SHAP values and prediction errors of POM semantic labels across test-set subtypes (y-axis). **(c)** Cumulative featureimportance of different models (y-axis) as a function of the number of features; the horizontal dashed line marks the 50% importance threshold; vertical dashed lines mark the number of features required to cross 50% importance threshold.

For the *D2Smell* team, features derived from pair model labels [7] contributed most strongly to predictions across the test-set, with a mean absolute SHAP value of 0.11 *±* 0.019. The *UMich CASI* team used 138 semantic labels from the POM model, grouped into 12 olfactory categories. The top three contributors were *“Fruity”, “Green & Herbal”*, and *“Nutty, Fatty & Roasted”*, with mean absolute SHAP values of 0.04 *±* 0.007, 0.03 *±* 0.006, and 0.03 *±* 0.005, respectively. The *ChemSenSim Lab* team utilized the full POM embedding, rather than probabilities, along with features such as predicted olfactory receptor responses, experiment type, and relative mixture size. SHAP analysis showed that the POM embedding contributed most to predictive performance (0.06 *±* 0.014). The *belfaction* team used 30 principle components of the

POM embedding and summarized mixture-level properties using statistical descriptors (mean, product, absolute difference, minimum, maximum, and U statistic). Among these, the absolute difference features provided the greatest contribution (0.10 *±* 0.004). The *PL21* team used a combination of chemical and semantic descriptors; among these, Dragon descriptors contributed most to predictions (0.081 *±* 0.040). Finally, team *Tamarin*’s predictions were driven primarily by a combination of Deepnose and Mordred features (0.06 *±* 0.042).

Among all feature types, POM-derived representations were the most widely used, appearing as probabilities (*UMich CASI*), full embeddings (*ChemSenSim Lab*), and dimension-reduced embeddings (*belfaction*). To further investigate their influence, we analyzed the relationship between SHAP values and prediction errors for the probability-based semantic labels used by *UMich CASI* (Figure 3b). Semantic labels showing statistically significant correlations between SHAP values and prediction errors are shown along the x-axis of Figure 3b. Positive Spearman correlations (red) indicate that higher SHAP values were associated with larger predictions errors, whereas negative correlations (blue) indicate that greater SHAP values were associated with smaller prediction errors. The labels *“violet”, “aromatic”* and *“natural”* most strongly increased prediction errors, whereas *“green”, “clean”*, and *“cognac”* were most strongly associated with decreased errors. Although these relationships are not straightforward to interpret, owing to the lack of experimental data directly linking semantic labels to perceptual mixture outcomes, they nonetheless provide an empirical basis for selecting informative POM features in future model development. In addition, this pattern cannot be explained by label frequency of the GoodScents and Leffingwell data used for training POM [3, 10] (see Figure S12).

To assess model compactness, we analyzed cumulative feature importance derived from SHAP values as a function of the number of features. Figure 3c compares how efficiently each model concentrates its explanatory power within its highest-ranked features (see Methods). Using the 50 % cumulative importance threshold (horizontal gray dashed line in Figure 3c), the *Tamarin, D2Smell*, and *UMich CASI* models were the most compact, achieving this threshold with fewer than 100 features. In contrast, the*ChemSenSim Lab* and *belfaction* models were less compact, requiring approximately 100 and 1000 features, respectively, to reach the same threshold. All models eventually plateaued to 100 %, though the least compact models (i.e., *ChemSenSim Lab* and *belfaction*) exhibit long tails of low-importance features that add little explanatory power.

Finally, to assess the stability of feature-importance ranking, we calculated Kendall’s coefficient of concordance (W) (see Methods). The resulting stability distributions are shown as ridgeline plots in Figure S12) for each test-set subtype and for all subtypes combined. Across models, W values peaked around 0.5, indicating moderate, but not strong, agreement in feature rankings. For the angle subtype, stability varied more widely across models, likely due to its smaller sample size (10 mixture pairs) relative to random (16 mixture pairs) and manual (20 mixture pairs) subtypes.

In the post-challenge phase, insights from the SHAP analysis, guided the development of a new model that integrated a unified feature set combining open-POM, Pair-model, Dragon, Mordred, and DeepNose features. Detailed specifications of this model are provided in Supplementary information and in Table S2.

### 2.4 Ensemble model maintains high predictive accuracy in novel validation set

In the post-challenge phase we developed a new validation set to evaluate both the test-retest reliability of experimental measurements and the performance of the ensemble model on an independent dataset. We first generated 10 base mixtures using a molecular co-occurrence probabilities which tries to mimic naturally occurring mixtures. Then we used POM predictions to generate non-overlapping target mixtures spanning 5 different distance quintiles. In total, 50 non-overlapping mixtures pairs were experimentally tested using direct odor similarity ratings (see Figure 4a and Methods). For calibration, we also measured similarity ratings for two single-molecule reference pairs: diallyl disulfide (garlic odor) versus allyl caproate (pineapple odor), and strawberry aldehyde (strawberry odor) compared with itself.

**Fig. 4.**
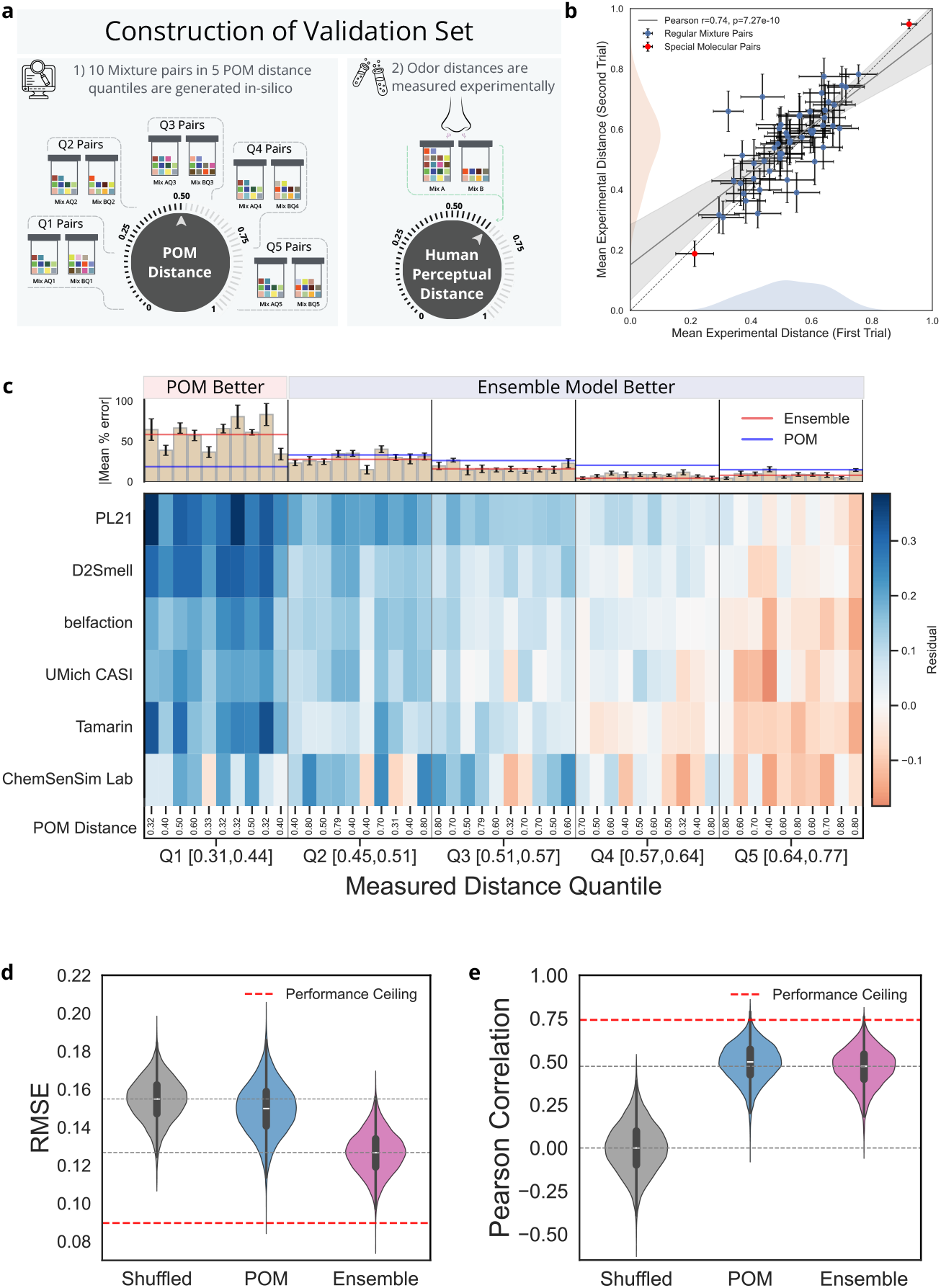
Construction of the validation set. **(a)** The in-silico mixture pair generations using POM in 5 distance quintiles (left), and experimental measurement of perceptual distance by human subjects (right), are shown. **(b)** The mean experimental distance values are shown for test (x-axis) and retest (y-axis) sessions using blue and red data-points. Blue data points correspond to 50 regular mixture pairs, and two red data points correspond to strawberry, garlic, and pineapple smelling molecule pairs; top right red point corresponds to garlic versus pineapple rating and bottom-left red point correspond to strawberry rating against itself, respectively. The error bars are standard errors of mean. Gray solid line shows the linear regression of regular mixture pairs with 95% confidence band. The light blue and light peach color distributions shows the distance distributions of test and retest sessions, respectively. **(c)** Residual heatmap showing team-experiment residuals, defined as the difference of model predictions and ground truth. Columns are grouped by measured distance quintiles; labels under the heatmap give each pair’s POM distance. Top bars plot mean absolute percent error along with standard error of means across teams per pair; two reference lines show quantile means for Ensemble (red) and POM (blue). **(d,e)** Distribution of RMSE **(d)** and Pearson correlation (y axis)**(e)** for POM and ensemble model (x axis), obtained via bootstrap sampling (n=10,000). The gray distribution represents chance-level performance from shuffled experimental distance values. The pink distribution represents ensemble model predictions. Gray dashed lines indicate median values for the ensemble model and shuffled baseline, and the red dashed lines are the performance ceilings obtained from test-retest reliability in two different trials.

Figure 4b shows the mean experimental distance values for the first (test, x-axis) and second (retest, y-axis) sessions. Blue points represent 50 mixture pairs, while the two reference molecular pairs are shown in red. The distributions of test and retest similarities are shown in light blue and light peach, respectively. A linear regression fitted to the regular mixtures pairs (blue points) is shown as a gray solid line with a 95% confidence band. Mean experimental distances were strongly correlated between test and retest sessions (Pearson’s r = 0.74, p-value=7.27e-10). The diallyl disulfide (Garlic smell) and allyl caproate (Pineapple smell) are judged far part, and Strawberry aldehyde (strawberry smell) compared to itself was judged very close, for both test and retest sessions. For regular mixture pairs, the distance measurements ranged in [0.29,0.76] interval for test and [0.31,0.78] interval for retest sessions, respectively.

We partitioned the measured perceptual distances into five quintiles, and computed model residuals for each mixture pair. Figure 4c shows a combined residual heatmap and absolute-error bar plot across all teams. Consistent with the test-set results, the validation-set exhibited a regression-to-the-mean pattern: models tend to overestimate perceptual similarity at lower distances and underestimate it at higher distances. The solid blue and red horizontal lines indicate the mean quintile errors for the POM and ensemble model, respectively. In the most similar quintile (Q1 [0.31,0.44]), the POM model showed lower error, whereas across the remaining quintiles (Q2-Q5 [0.45,0.77]), the ensemble model consistently outperformed POM with smaller errors in each quintile.

To evaluate the ensemble model’s performance on the validation set, we performed bootstrap sampling (n=10,000) to estimate the distribution of RMSE and Pearson correlation for both the POM and ensemble models (see Methods). We also established a chance-level baseline by randomly shuffling the experimental distance values. As shown in Figure 4d,e) both models outperformed the shuffled baseline. The ensemble model achieved the lowest RMSE (see Figure 4d), while both models exhibited comparable Pearson correlations (Figure 4e). No model reached the performance ceiling defined by the upper bound of the measurement error, as determined by the test-retest correlation.

## 3 Discussion

Prior to the training of advanced machine learning (ML) models made possible by the generation of the necessary data, predicting the odor perception of single molecules from their molecular structure had long remained a notoriously difficult problem [13]. This challenge has been addressed in the last decade, as demonstrated through the first DREAM olfaction challenge [1] and the development of the Principal Odor Map (POM) [3]. However, achieving high-fidelity prediction of olfactory mixture similarities remained an open problem. In this work, we leveraged a large-scale community effort, through the second DREAM Olfaction Challenge, a global collaborative effort to improve the prediction of olfactory mixture similarity. We unified six heterogeneous psychophysical datasets from three separate studies to create a comprehensive training dataset. Twenty-six international teams independently developed ML models to predict odor similarity, optimizing for low RMSE and high Pearson correlation on a hidden test-set of 46 mixture pairs. The competition resulted in a four-way tie among top-performing models. Following the challenge, an independent validation set of 50 olfactory mixtures was developed to estimate prospective model reliability.

From the 19 submitted models, we analyzed the six top-performing submissions, all of which achieved RMSE < 0.1 and provided fully reproducible and open-source code. The competition resulted in a four-way tie among top performers (*D2Smell, UMich CASI, ChemSenSim Lab*, and *belfaction*). This tie likely reflects both the relatively small size of the test set (46 mixture pairs) and the fact that it was drawn from a different distribution than the training data, making fine discrimination among similar high-performing models difficult. The tie does not represent a performance ceiling, as split-half reliability measurements indicate further room for improvement. Averaging predictions from the six top models produced an ensemble model with a median RMSE of 0.08 and a Pearson correlation of 0.57 on the test set. The ensemble outperformed each individual model as well as four SOTA baselines: the Snitz angle model, the POM model, the semantic model, and the aroma-chemical pair model. On the prospective validation set, the ensemble maintained strong performance, with RMSE of 0.13 and achieved comparable correlation to the POM model. The ensemble’s superior performance indicates that combining diverse weak learners can better approximate the underlying latent perceptual manifold while reducing noise, consistent with the advantages of ensemble learning observed in computer-vision, natural language processing and many other domains [14–16].

Most individual models, and therefore the ensemble, performed best on the manual and angle subtypes of the test-set, particularly in reducing prediction error. This suggests that the top models captured higher-order perceptual structure of systematically designed olfactory mixture pairs compared to randomly generated mixtures.

Despite the wide range of modeling strategies and feature sets across teams, SHAP analysis revealed that semantic descriptors as well as semantic features derived from the POM and the pair model consistently provided the most predictive information. POM features were used in several forms, including semantic label probabilities, embeddings, and dimension-reduced representations, all of which contributed disproportionately to model accuracy. Semantic labels from the pair model also made substantial contributions, particularly in the *D2Smell* model. Together, these results suggest that although POM was trained exclusively on single molecules, the perceptual distance between olfactory mixtures can also be effectively represented in a low-dimensional semantic space. In other words, predicting mixture perception from component molecules may not be as challenging as previously theorized [17]. This challenges the notion that mixture perception involves fundamentally different mechanisms than single-molecule perception and suggests that the primary challenge lies in the initial structure-to-percept mapping rather than the component-to-mixture transformation.

Prior studies have shown that POM effectively captures olfactory relationships across varied datasets, outperforming generic molecular descriptors [18]. Our results extend this finding to mixture perception and suggest that semantic representations provide the most efficient features for predicting olfactory perceptual similarity. In terms of model compactness, three of the most accurate individual models reached 50% cumulative SHAP importance using fewer than 100 features, indicating that accurate predictions can be achieved with compact feature sets (Figure 3c).

Despite the ensemble’s strong performance, a gap remains between model predictions and human test-retest reliability (r=0.74 on validation set, r=0.70 split-half reliability on test set). This gap may reflect the inherent difficulty of diagnosing mixture similarity errors. Error in predicting mixture similarity could arise at multiple levels: (1) the structure-to-percept mapping for individual components, (2) the component-to-mixture integration, or (3) the mixture-percept-to-similarity metric. Disentangling these error sources is challenging with current data, as we lack systematic measurements of component percepts, predicted mixture percepts, and similarity judgments within the same experimental framework.

Across all models and test set subtypes, relative error showed a negative correlation with experimental distance values, indicating that models tend to bias predictions toward intermediate values—overestimating similarity at lower perceptual distances and underestimating it at higher ones. This regression-to-the-mean pattern reflects limited training data at the distributional extremes (1e). Prediction error is lowest for mixtures rated in the midrange (0.5 to 0.8), where training data are most abundant, and increases for distances below 0.5 or above 0.8.

Several limitations constrain current model performance and generalization. First, although we unified multiple measurements paradigms, including triangle tests and similarity ratings, by mapping all measurements onto a 0 to 1 distance scale, this approach overlooks paradigm-specific biases [19] and residual noise arising from different administration techniques (*e*.*g*, headspace via pads in jar [5] vs. liquid in vials [4, 9]).

Second, the training data are unevenly distributed across the similarity scale, with higher density around moderate distances (0.5 to 0.8). This imbalance occurs because, in our experience, randomly generated mixtures tend to fall within this range. Without a pre-existing model to guide mixture design, it is difficult to generate sufficient mixture pairs at the extremes of the distribution. At low perceptual distances, all models overestimated the perceptual distance, as shown in Figure S10. Improving performance at the distributional extremes will require actively designing and measuring new mixture pairs.

Third, most datasets used here include only intensity-matched components, with the exception of the Ravia dataset. Odor quality can shift with intensity [20–22], and, within mixtures, more intense components can mask weaker ones [23]. Since real-world odors rarely consist of intensity-balanced components, future models must account for intensity to improve their generalization to natural odors.

Fourth, our post-challenge effort to develop a refined model using only the highest-contributing features (open-POM logits, pair-model embeddings, Dragon and Mordred descriptors, and DeepNose features) achieved performance comparable to individual top performing teams rather than the ensemble (see Figure S13). This suggests that while feature selection and model architecture are important for learning olfactory perceptual similarity, achieving ensemble-level performance requires the diversity of modeling approaches rather than simply identifying optimal features. The post-challenge model’s performance may also reflect fundamental model limitations when training data are scarce.

Finally, both the test set and validation set use molecules from a limited palette and whether models can generalize to completely novel molecules not represented in training remains an open question that will require future evaluation.

Closing the gap between model performance and human reliability will require several advances. Most critically, we need larger, more systematically designed datasets that span the full range of perceptual distances with more specific perceptual information and intensity variation. Ideally, future datasets would include measurements at multiple levels—component percepts, mixture percepts, and pairwise similarity judgments—enabling more precise diagnosis of error sources.

Our results provide a high-fidelity, machine-readable framework for compressing high-dimensional chemical and semantic information into a one-dimensional perceptual metric. This metric enables efficient *in-silico* experiments that were previously impractical or computationally expensive. For example, virtual olfactory metamers could now potentially be generated efficiently and tested experimentally. Accurate and robust mixture-similarity metrics are foundational for a wide range of technological applications, including content-aware compression for digital olfaction, non-invasive health monitoring [24, 25], generative fragrance design [26–28], next generation electronic nose sensors [29, 30], and development of olfactory foundation models [31].

We encourage the community to expand publicly available psychophysical datasets, along with metadata on concentration and measurement techniques. Expanding these olfactory datasets is the most direct path to training more robust, generalizable models capable of learning the underlying structure of olfactory mixture perception and bringing us closer to understanding olfaction.

## 4 Methods

### 4.1 Training Dataset

Training dataset composed of six datasets of odor-similarity measurements from three different studies [4, 5, 9] (see Table S1). Each dataset is originally designed with different objectives. For instance, the Snitz dataset [4] focused on developing an odor-similarity metric between mixture pairs based on chemical descriptors, whereas the Ravia dataset [5] was designed to create olfactory metamers, *i*.*e*. non-overlapping mixture pairs with similar olfactory perception. In contrast, the Bushdid dataset [9] was designed to measure human olfactory discrimination capacity by increasing the overlap of mixture pairs under comparison. Because these datasets were designed for different goals, they vary in mixture composition and measurement techniques [32], with some using triangle tests, while others leverage similarity ratings (see Figure S5). For the DREAM Challenge, we standardized and merged available measurements into a unified dataset, comprising 168 unique mono-molecules, 731 unique mixtures, and 507 mixture pairs measurements. Mixture pair distances were mapped onto a continuous perceptual scale from 0 (indistinguishable) to 1 (maximally distinct).

### 4.2 Development of the test-set

A total of 34 participants (20 females) aged between 18 and 50 years (median age: 40 years), were recruited at The Rockefeller University. All subjects provided written informed consent to approved procedures by the Rockefeller University institutional review board (IRB: LVO-0857). Participant were excluded if they reported abnormal olfactory function, upper respiratory illness, or allergies to fragrances.

A total of 92 unique olfactory mixtures which form 46 mixture pairs were used. Each mixture contained 10 non-overlapping odorous components selected from previously intensity matched set of molecules [9]. The mixture pairs were classified into three subtypes: i) random (32 mixtures generated randomly), ii) manual (40 mixtures designed based on primary semantic descriptors), and iii) angle (20 mixtures generated based on Snitz angle methodology [4])

Participants performed triangle test, three-alternative forced-choice discrimination tasks in which they were asked to identify the odd smelling odor vial from three presented vials. Each triangle test consisted of three vials. Two vials contained the same mixture. The third vial contained a different mixture. To assure that the discrimination is based on differences in odor quality and that subjects do not merely use slight differences in intensity to distinguish the mixtures, all three vials were presented at different dilutions (undiluted, 1:2 dilution, 1:4 dilution). Vials were presented in random order, and instructions and collection of responses were administered via a computerized interface. To avoid adaptation and olfactory fatigue, we enforced a 90 second rest period before subjects could move from one three-alternative forced choice task to the next. For each mixture pair, a total of 136 or 140 measurements were collected. All measurements were conducted between October 29, 2014 and March 3, 2015.

### 4.3 Competition rules

1. Participants had to register for a Synapse account at synapse.org and become certified users. Certification ensured that all participants had completed the required training on data use and responsible research conduct, as Synapse hosted the challenge data.
2. Teams were allowed to add additional experimental data to the original challenge training data.
3. The use of external dataset as model features for training was permitted.
4. Augmenting the provided training data and employing ensemble model was allowed.
5. Teams could submit up to three predictions on the held-out leaderboard (LB) dataset containing 46 measurements, to assess their model performance.
6. Each team submitted a single prediction on the hidden test-set, also consisting of 46 measurements.
7. Reproducibility was required for all submissions. Participants were required to provide code and detailed instructions for replicating their models. The top-performing models were checked for reproducibility by organizers.
8. To qualify for consortium-level authorship, a write-up must be submitted. The write-up should include detailed information on the methodologies, external data sources used (if any), and accompanying source codes.

In total, 26 teams submitted 159 Leaderboard predictions, and 19 teams submitted test-set predictions. Detailed information on the competition and its guidelines can be found on the DREAM Challenge webpage (https://www.synapse.org/Synapse:syn53470621/wiki/). The detailed implementation of all models is provided in the Supplementary information.

### 4.4 Performance evaluation metric

We calculated root mean squared error (RMSE) and Pearson correlation to evaluate olfactory distance prediction performance. By integrating these two metrics, we provided a comprehensive evaluation of each model’s accuracy and predictive power, to ensure that both the magnitude of the prediction errors and the consistency of the predicted trends were taken into account. Throughout the paper, we define a chance-level baseline by randomly shuffling the experimental distance values in both test and validation sets.

### 4.5 Bootstrapping analysis of model performance

To determine top-performing teams, we calculated the Bayes Factor to quantify how frequently each team ranked first in both RMSE and Pearson correlation, across 10,000 bootstrapped simulations. In each iteration, 10% of the test was randomly added to introduce variability. The competition resulted in a four-way tie among *D2Smell, UMich CASI, ChemSenSim Lab*, and *belfaction* teams.

### 4.6 State-of-the-art (SOTA) models

We implemented SOTA benchmarks using, linear, logarithmic, or lasso regression methods. Specifically, we evaluated four established models: Snitz angle [4, 5], open-POM (an open source implementation of POM [3, 10]), a semantic model [6], and an aroma chemical pair model [7]. For the Snitz angle model, we computed the Snitz angle between all mixture pairs in the training dataset using 21 Dragon descriptors [4]. We then performed linear and logarithmic regression to identify the best-fitting relationship between Snitz angle and experimental distance values of training set. The best fit is eventually used to predict the test-set olfactory perceptual distance. For POM, we first generated 138 dimensional semantic descriptor probabilities for each molecule individually. Each mixture was then represented by averaging the semantic descriptor probabilities of its constituent molecules, leading to a 138 dimensional vector per mixture. From these vector representation, we calculated distance metrics from verity of quantities, including Pearson correlation, cosine similarity, Euclidean distance, and angle for all mixture pairs. As with the Snitz angle approach, we applied linear and logarithmic regressions to these metrics in training set and used the best fit as a predictive model to obtain olfactory perceptual distance for test-set. For the semantic model [6], predictions were directly generated from an open-source implementation based on lasso regression. Finally, for aroma chemical pair model [7], we adopted the same computational pipeline as the POM model, but used the mixture-level probabilities derived from pair model descriptors (see Figure S8)

### 4.7 Feature importance analysis using SHAP values

We used SHapley Additive exPlanation (SHAP) values [11, 12] to analyze the importance of different classes of features.

To assess model compactness, features were first sorted by their mean absolute SHAP values, and cumulative sums were computed to determine each feature’s contribution relative to total feature importance. The slope of this cumulative curve provides a measure of model compactness, indicating how many features are needed to reach a given explanatory threshold.

To calculate Kendall’s coefficient of concordance (W), test-set predictions were divided into two consecutive, equally sized blocks of mixture pairs. Within each block, features were ranked according to their SHAP values, and Kendall’s W was then calculated across blocks to quantify rank stability for each feature.

### 4.8 Development of the validation set

#### 4.8.1 Generation of validation mixture pairs

To approximate real-world odor mixtures, we first aimed to quantify molecular cooccurrence in naturally occurring odorants found in food items, essential oils, and other complex mixtures. We assembled a natural occurrence dataset for 97 test-set molecules from three sources: the Leibniz-LSB@TUM Odorant database [33], the EOU database [34], and the Good Scents database [35]. We then computed the pairwise co-occurrence probabilities among these molecules. Using this dataset, probability of semantic labels from POM, and greedy search, we generated two classes of in-silico mixtures (Class A and B):

- **Class A Mixtures:** We randomly selected five molecules from the 97 test-set molecules, then we added five additional, non-overlapping molecules with the highest co-occurrence probabilities. Using this procedure, we generated 100,000 unique Class A mixtures, each containing 10 molecules. From this pool, we selected a set of 10 mixtures with minimal overlap in their components, Figure S14. The Class A mixtures served as the baseline reference set.
- **Class B Mixtures:** We implemented a greedy search algorithm using the open-POM [3], and the same palette of 97 test-set molecules in our greedy search. We selected non-overlapping molecule based on a predefined Pearson correlation to each Class A mixture. For each Class A reference, mixtures were generated at 10 correlation levels (quintiles: 0, 0.1, 0.2,…,1.0) through the greedy search. This process resulted in some duplicates due to challenge of generating mixtures in the extremes of similarity or dissimilarity in POM space. After removing duplicates, we retained 66 unique Class B Mixtures, each consisting of 9 to 11 components. The Class B mixtures represent perceptually tuned distances from the baseline Class A mixtures, based on POM embeddings.

We finally used Class A and B as mixture pairs to measure olfactory perceptual similarity of 50 pairs.

### 4.8.2 Olfactory similarity measurements of the validation set

A total of 16 participants (7 females) aged between 18 and 55 years (mean age: 32 years), were recruited at the Monell Chemical Senses Center. All subjects provided written informed consent to approved procedures by the University of Pennsylvania institutional review board (IRB818208). Participant were excluded if they reported abnormal olfactory function, upper respiratory illness, or allergies to fragrances.

A total of 59 unique odor mixtures which form 50 mixture pairs were used. Each mixture contained 9 to 11 non-overlapping odorous components, selected from a set of 94 single molecules that had been intensity-matched. Twenty-one of them were selected from a previously intensity matched set of molecules [9, 36]. Seventy-three of them were intensity matched during the development of validation set at the Monell Chemical Senses Center, using method described previously [4, 9].

Participants were first trained using the Napping® technique [37], where they arranged odor samples on a two-dimensional space by placing similar odors close together and dissimilar ones farther apart. They also learned to rate the similarity between odor mixtures on a 0–100 scale, with 0 indicating “highly similar” and 100 “not at all similar.” To validate their understanding, participants evaluated 40 predefined mixture pairs, with key reference points including a highly similar strawberry–strawberry pair and a highly dissimilar pineapple–garlic pair. In the actual test, in addition to the 50 target mixture pairs, two reference pairs were included: one composed of a mixture and its identical replicate to serve as a high-similarity reference, and another consisting of a garlic–pineapple pair to serve as a low-similarity reference. Participants rated the perceptual similarity of odor mixture pairs using a direct similarity rating task conducted in two sessions spaced at least one day apart. The first session for test, and the second session for retest. On each trial, two vials were presented in random order, and participants were asked to sniff both vials and record their similarity rating, ranging from identical to extremely different, via a computerized interface. For each mixture pair, a total of 30 measurements were collected. All measurements were conducted between July 8 and July 29 of 2025.

## Supplementary information

The online version contains supplementary material available at:

## Declarations

### Data and Code availability

All datasets, models, evaluation scripts, and figures are available at the following repository https://github.com/Satarifard/DREAM-olfactory-mixtures-prediction-challenge/

### Author contribution

P.M and J.M designed the DREAM Challenge. J.M., P.M, J.A., G.A., organized the DREAM Challenge. A.K. and L.B.V. provided scientific advice to challenge organizers. The top-performing four models were designed by the following teams: D2Smell (V.S., W.Y., M.B., D.K., N.A.C., N.S., A.R.), UMich CASI (Y.H., Z.W., S.Y., A.T.), ChemSenSim Lab (M.H., M.L., J.T.), and belfaction (P.I., R.D., A.G., J.B., F.K.N., A.G., C.V.). The remaining models were proposed by the DREAM Olfactory Mixtures Prediction Consortium. V.S., Y.H., P.I., M.H., and M.L. performed SHAP value analysis. V.S. conducted all post-processing, including SOTA analysis and generated figures. V.S., P.M., J.M. interpreted the results of the challenge and all follow-up analyses. V.S., P.M., and J.M. wrote the paper with input from all other authors. L.S. developed the post-challenge model. V.S. and L.S. developed in-silico validation set mixture pairs. A.K. and X.S. collected the experimental distance values for the test set and validation set, respectively. B.S.L., A.K., L.B.V., S.F., A.T., J.T., C.V., M.B., D.K., N.S., N.A.C. supervised the research.

