## Supplementary Information for "High-Fidelity Tuning of Olfactory Mixture Distances in the Perceptual Space of Smell Through a Community Effort"

[1]Human Nature Lab, Yale University, New Haven, CT, USA.
[2]Department of Computer Science, Boston University, Boston, MA, USA.
[3]Department of Statistics, University of Michigan, Ann Arbor, MI, USA.
[4]Department of Public Health and Primary Care, KU Leuven, Kortrijk, Belgium.
[5]itec, imec research group at KU Leuven, Kortrijk, Belgium.
[6]Institut de Chimie de Nice, Université Côte d'Azur, CNRS, Nice, France.
[7]Department of Computer Science, University of Oxford, Oxford, United Kingdom.
[8]Monell Chemical Senses Center, Philadelphia, PA, USA.
[9]School of Electrical Engineering and Computer Science, KTH Royal Institute of Technology, Stockholm, Sweden.
[10]Cornell Tech, Cornell University, New York, NY, USA.
[11]Cold Spring Harbor Laboratory, Cold Spring Harbor, NY, USA.
[12]Sage Bionetworks, Seattle, WA, USA.

[13]Department of Chemical Engineering and Applied Chemistry, University of Toronto, Toronto, Canada.
[14]Laboratory of Neurogenetics and Behavior, The Rockefeller University, New York, NY, USA.
[15]Howard Hughes Medical Institute, The Rockefeller University, New York, NY, USA.
[16]Department of Neurobiology, Weizmann Institute of Science, Rehovot, Israel.
[17]Department of Neuroscience, University of Pennsylvania, Philadelphia, PA, USA.
[18]Health Care and Life Sciences, IBM Research, New York, NY, USA.
‡The list of consortium authors and affiliations is provided at the end of this document..

†These authors contributed equally to this work.

### 1 Model Details

Computational details for the top six models used to construct the ensemble are presented below, along with the excluded models and the post-challenge model. The model descriptions are slightly edited version of write-ups submitted to the DREAM Olfactory Mixtures Prediction challenge (https://www.synapse.org/Synapse:syn53470621/wiki/)

#### 1.1 D2Smell

The team used a chemical-aroma pair model that is trained on experimental measurements of semantic labels of binary mixtures [2]. By combining these sets of single molecule and binary mixture semantic labels along with the compound identifiers (CID), they trained a machine learning model to predict the olfactory mixture similarity. Team used XGBoost as the machine learning technique that has been shown to be advantageous for smaller datasets such as the olfactory mixture dataset. Regarding feature selection, they used 21 Dragon chemical descriptors that have been shown to be more relevant to the olfactory mixture [3, 4]. Team utilized these chemical descriptors to train a model for predicting 19 semantic labels as well as the intensity and pleasantness of single molecules [5, 6]. Additionally, they employed an aroma-chemical pair model to predict binary mixture olfactory semantic descriptors [2]. Moreover, team found that the quantile transformer, along with data augmentation, improves the performance of the predictive model on the leaderboard set results.

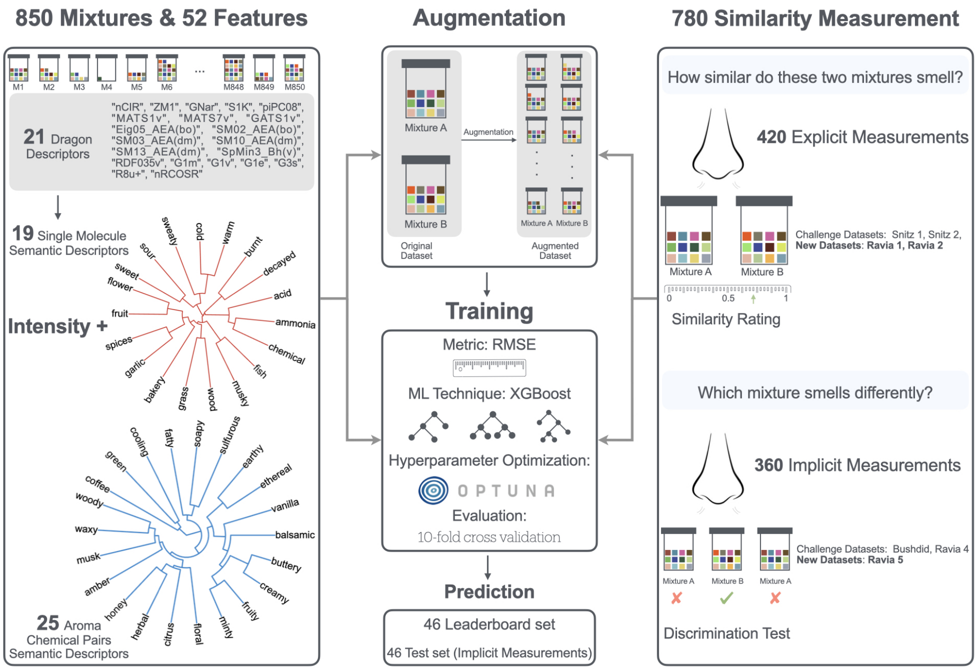


**Fig. S1** This schematic workflow illustrates the complete workflow involved in the study of D2Smell team, including model input, feature selection, data augmentation, and model training.The hierarchical clustering of semantic descriptors is conducted based on the cosine similarity of word embeddings derived from the Word2Vec model [1].

##### 1.1.1 Methods

The schematic workflow used in the study conducted by D2Smell team is summarized and shown in Figure S1. In the following the details of different steps leading to the final predictions will be discussed.

**Dataset Curation**. The original dataset provided by the DREAM challenge organizers [7] included four different datasets from three separate studies: Snitz 1, Snitz 2, Ravia 4, and Bushdid. In the early stages of model training, team discovered several parsing errors in the Snitz 1 and Snitz 2 datasets, as well as numerous mislabeling issues and missing CIDs in the Bushdid dataset. Moreover, they realized the possibility of adding three more experiments to the original dataset that could potentially improve the model's performance. In light of these points, the team curated a new dataset from original publications [3, 4, 8] and added three more datasets: Ravia 1, Ravia 2, and Ravia 5, to the full training dataset. They ultimately compiled a dataset consisting of 850 unique mixtures and 780 similarity measurements.

**Feature Selection.** Inspired by previous studies [3, 4], they utilized 21 Dragon chemical descriptors, including 'nCIR', 'ZM1', 'GNar', 'S1K', 'piPC08', 'MATS1v', 'MATS7v', 'GATS1v', 'Eig05AEA(bo)', 'SM02AEA(bo)', 'SM03AEA(dm)', 'SM10AEA(dm)', 'SM13AEA(dm)', 'SpMin3Bh(v)', 'RDF035v', 'G1m', 'G1v', 'G1e',

'G3s', 'R8u+', and 'nRCOSR', in conjunction with an XGBoost regressor to predict pleasantness, intensity, and 19 semantic descriptors: 'bakery', 'sweet', 'fruit', 'fish', 'garlic', 'spices', 'cold', 'sour', 'burnt', 'acid', 'warm', 'musky', 'sweaty', 'ammonia', 'decayed', 'wood', 'grass', 'flower', 'chemical' for each molecule. Team then trained a model with 33 aroma-chemical pair semantic descriptors [2], which included: 'alliaceous', 'coffee', 'floral', 'fruity', 'green', 'herbal', 'minty', 'sulfurous', 'waxy', 'balsamic', 'aldehydic', 'buttery', 'caramellic', 'creamy', 'earthy', 'ethereal', 'fatty', 'fermented', 'musk', 'soapy', 'spicy', 'tropical', 'woody', 'amber', 'cooling', 'citrus', 'animal', 'berry', 'honey', 'vanilla', 'nutty', 'musty', 'camphoreous'. They dropped 8 labels 'aldehydic', 'fermented', 'spicy', 'tropical', 'animal', 'berry', 'Nutty' and 'musty' due to their low performance with an AUROC below 0.5, while all other labels showed good performance with average AUROC of 0.80. For each mixture, they averaged over the predictions of semantic descriptors for all possible binary combinations of compounds in the mixture. Finally, each mixture was represented by a binary vector of molecular IDs (235 unique molecules), combined with intensity and the highest single molecule semantic descriptor values, as well as 25-dimensional binary mixture semantic descriptors. This set of features leverages both single molecule and binary mixture semantic descriptors, along with molecular overlap of mixtures, to enhance the prediction accuracy and interpretability of the model.

**Data Augmentation.** To address the challenge of model overfitting due to a limited dataset, they implemented a data augmentation strategy by manipulating molecular counts in mixtures. This involved either increasing or decreasing the experimental similarity by a fixed amount. Specifically, if a molecule present in mixture A was added to mixture B, which lacked it, the experimental distance between the two mixtures decreased by a predetermined amount, a parameter fine-tuned through hyperparameter tuning. This adjustment resulted in significant data expansion: from an initial count of 780 similarity measurements (treating A-B and B-A as distinct pairs), the dataset grew to 4,078 synthetic similarity entries with the addition of one molecule per mixture, and to 7,331 entries with two molecular adjustments. They further augmented the data by excluding common molecules; specifically, if mixture A and mixture B shared a molecule, removing it from both increased their experimental distance by a predetermined amount. This step increased the total dataset size to 10,692, of which approximately 93% was synthetic data. Though not fully optimized, preliminary tests indicated that these augmentation steps, adding up to two molecules and removing one, improved the model performance on the leaderboard set.

**Model Training and Prediction.** For model training, they utilized XGBoost as the primary machine learning (ML) technique to train predictive models for olfactory mixture similarity. The fast training speed of this technique is well-suited for small datasets and requires fewer computational resources. Built-in regularization helps prevent overfitting, which is crucial for handling small data sets. The team employed root mean squared error (RMSE) as the metric for calculating the loss function and used Optuna for hyperparameter optimization. Furthermore, they implemented 10-fold cross-validation to evaluate the model's performance. Finally, they used the trained model to predict the outcomes on the test set.

##### 1.1.2 Summary

In summary, D2Smell team have utilized XGBoost to train a predictive ML model for olfactory mixture similarity. To address the limited size of the experimental data, they implemented data augmentation and various regularization techniques, along with 10-fold cross-validation to prevent overfitting. Their model demonstrates excellent performance on the full dataset (RMSE=0.08 and Pearson Correlation=0.89) on each separate dataset, as well as on the leaderboard dataset. In addition to the discussed efforts, they experimented with using single molecular semantic descriptors from open-POM [9, 10] as input features, in combination with symbolic regression, random forest, elastic net, and lasso as ML techniques. However, none of these efforts resulted in improved performance particularly for correlations. The source code of D2Smell model is available at: https://github.com/Satarifard/CWYK-Olfboost.

#### 1.2 ChemSenSim Lab

Mammals perceive and interpret a myriad of olfactory stimuli through a sophisticated coding mechanism involving interactions between odorant molecules and hundreds of olfactory receptors (ORs). These interactions generate unique patterns of activated receptors, forming what is known as the combinatorial code, which the brain interprets as distinct smells. Olfactory input is thus conveyed by odorant molecules and encoded through this combinatorial code. To solve the odor discriminability task, they propose to use the recent advances in OR activity and odor quality predictions. Utilizing state-of-the-art methodologies by Hladiš et al. [11] and Lee et al.[9], they combine predicted combinatorial code and Principal Odor Map (POM) embeddings, respectively, as the representation of the components.

Mixtures introduce synergistic and antagonistic effects leading to a non-trivial function combining the properties of the components into the properties of the mixture. Thus, they introduce a learned aggregation function to combine the diverse information within odor mixtures' components.

Traditional distance functions, such as Euclidean distance, fall short in capturing the intricate interactions in olfactory perception. Therefore, they incorporate a metric learning approach based on Siamese networks on top of the aggregation function to better capture the nuanced relationships between different odorant combinations. Additionally, as demonstrated by Bushdid et al. [8] and Weiss et al.[12], discriminability depends on the mixture size, which they also use as an input. Their methodology thus allows for more accurate and reliable discrimination of olfactory mixtures, overcoming the limitations of classical distance metrics.

##### 1.2.1 Methods

**Data Pre-processing.** Data were gathered from the challenge data repository and manually curated to correct or add missing CIDs. Special attention was given to isomers, as human olfaction can discriminate between them. Additionally, missing discrimination pairs from the studies by Bushdid et al.[8], Ravia et al.[4], and Snitz et al.[3] were collected. Using these three data sources, various discriminability experiments were conducted. Each pair was labeled according to the type of experiment:

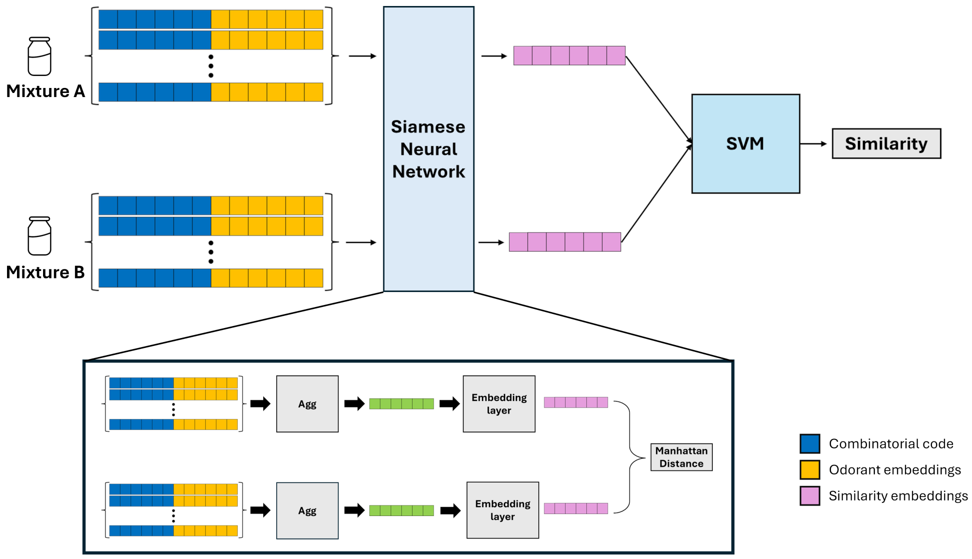


**Fig. S2** This schematic workflow illustrates the complete workflow involved in ChemSenSim Lab model.

"Triangle" for triangular tests, "1 - Similarity" for explicit similarity experiments, and "Same-Different" for same-different tasks.

**Encoding and Representation of Odor Mixtures.** Each molecule within a mixture was encoded using its combinatorial code and its embedding from an odor quality prediction task. The combinatorial code was generated using the procedure by Hladiš et al.[11] trained on the M2OR dataset [13], predicting the probability of activation for 385 reference olfactory receptors as defined in the RefSeq [14], resulting in a 385-dimensional vector per molecule. Additionally, team concatenated this vector with the 256-dimensional embedding of the odorant molecule derived from the odor quality prediction model of Lee et al.[9, 10], yielding a combined vector of size 641.

To account for the various types of experiments, team performed one-hot encoding, adding a 3-dimensional vector. The ratio of the sizes of the two mixtures

$$\frac{\min(|M1|, |M2|)}{\max(|M1|, |M2|)}$$

where $|M1|$ and $|M2|$ are numbers of components in mixtures 1 and 2, respectively, conveying size information, was also included, resulting in a final vector size of 645 per molecule. Each mixture was then represented by an n×645 matrix, where n is the number of molecules within the mixture. During the model training, each mixture was padded, resulting in a final matrix of 61×645.

**Siamese Neural Network Aggregation Function.** The aggregation function began with a fully connected layer, followed by layer normalization [15] and a dropout [16] with a 0.3 probability to prevent overfitting. An attention mechanism was applied to weigh different components of the mixture, with padding masks considered to handle variable-length inputs.

**Siamese Network.** The Siamese network consisted of the aggregation function followed by three sequentially arranged fully connected layers. The first layer transformed the input dimension to 1024, followed by batch normalization and a dropout layer. The second layer reduced the dimension to 512, again followed by batch normalization and dropout. The third layer further reduced the dimension to 256, with batch normalization and dropout applied similarly. The network independently processed two input mixtures, each passing through the same sequence of transformations to generate embeddings. These final processed embeddings are used to calculate the Manhattan distance normalized by the input dimension. Mean squared error (MSE) loss function was used to compare the predicted and true distances.

**Training Procedure.** The model was trained using an Adam optimizer with a learning rate of 0.001, running for 100 epochs with a batch size of 300. Each training iteration involved the following steps: (1) Passing mixture pairs and their padding masks through the aggregation function to produce aggregated outputs. (2) Processing these outputs through the rest of the Siamese network to generate embeddings. (3) Computing the Manhattan distance between embeddings and comparing it to the true distances using the MSE loss function. (4) Updating parameters accordingly.

**Post-processing using SVM.** Finally, in the last stage of the procedure, the learned embeddings from the Siamese network were used as an input to a Support Vector Regression (SVR) model. This model was then trained on the same dataset and its predictions were submitted to the challenge.

##### 1.2.2 Summary

In this study, ChemSenSim Lab team developed a novel deep metric learning approach to address the challenging task of olfactory mixture discriminability. By leveraging the strengths of Graph Neural Network (GNN) embedding and the combinatorial code of olfaction, their method effectively captured the complex interactions between odorant molecules and olfactory receptors. The integration of a Siamese network allowed them to exploit the nuanced relationships between different odorant combinations, surpassing the limitations of traditional distance metrics like Euclidean distance. Their results demonstrated that the model reliably discriminated between olfactory mixtures, effectively accounting for various mixture sizes. The source code of ChemSenSim Lab model is available at: https://github.com/chemosim-lab/ChemoDREAM.

#### 1.3 belfaction

Throughout this challenge the belfaction team sought to build on this existing work and create predictive models which leverage both objective molecular properties like physio-chemical descriptors, Morgan fingerprints and complex molecular structures, as well perceptual properties relating to the perceived smell of mixtures across 19 semantic descriptors [5]. They experimented with a number of ways of incorporating this information, as well as the methodological innovations in [9, 17, 18], and found that Extremely Randomized Trees applied on diverse statistical descriptors of mixture pairs performed best.

##### 1.3.1 Methods

In this section, team first present a number of methods explored to obtain numeric descriptors of molecules. They then describe the submitted models throughout this challenge.

**Molecule perception embeddings.** To generate perception embeddings for molecules, team used 19 odor descriptors from 338 molecules previously utilized in the DREAM Olfaction Prediction Challenge [5]. The input descriptors were in the form of Simplified Molecular-Input Line-Entry System (SMILES) strings. For this task, they employed the Chemprop package [17], which leverages directed message-passing neural networks (D-MPNNs) to predict various molecular properties by capturing complex molecular features. To obtain the perception embeddings, they extracted the activations of the last layer of the trained model. While the default parameters were used, they made specific adjustments: dropout was set to 0.5 and the message hidden dimension was set to 30 to mitigate overfitting. The feed-forward network hidden dimension was adjusted to 128 to achieve an embedding size of 128.

**Molecule perception predictions.** The team considered a number of approaches and data sources to generate molecule level perception predictions for the 19 descriptors from the 2017 DREAM challenge dataset. The best performing models combined physio-chemical descriptors like in [5] and [4] for each molecule with molecular structure information from morgan fingerprints and the embeddings obtained by applying D-MPNNs to the SMILES representations from the previous step. Each descriptor was modeled independently using the LightGBM implementation of tree boosting and tuned using bayesian hyperparameter optimization. These models improved on the performance of [4, 5] on the individual molecule perception level, and while not selected as our final mode for mixture similarity predictions, they improved on [4] when averaging mixture perceptions based on the component molecule predictions and including these differences as features to the baseline model described below.

**SMILES-BERT embeddings.** Also using SMILES string as input, they extracted embeddings using a variant of SMILES-BERT [17], a deep learning architecture composed of 12 transformer autoencoder layers with 12 attention heads. In the intermediate layers, the Gaussian error linear unit (GELU) activation was employed and the dropout rate was set to 0.1, whereas the hyperbolic tangent was used as the activation function in the pooling layer. Furthermore, the network was trained using molecules from the ChEMBL database. They extracted 178 embeddings, i.e., activations of the last layer, which reflect the general structure of the molecule.

**Hypergraph-based features.** They used a hypergraph to model the complex relations among different elements. A hypergraph is an extension of a graph where a hyperedge can connect more than two vertices (nodes) [19]. They treated mixtures, molecules, and perceptions as nodes in the hypergraph. They defined three types of hyperedges: hyperedges that link a mixture to its constituent molecules, hyperedges that link each molecule to its perceptions, and hyperedges that link similar mixtures based on a given distance. By constructing such a hypergraph with the specified nodes and hyperedges, they could measure the graph closeness between any two mixtures in the hypergraph. This graph closeness can serve as an additional feature for mixture pairs.

**Tree-embeddings.** They leverage the structure of each tree in a decision forest to characterize both mixtures and molecules [20]. The initial step is to build a Random Forest on a pretext task. They experimented with several input/target options to define pretext tasks. For example, binary vectors representing each mixture which indicate the presence of each molecule of our dataset could be used as inputs. As output, they could average the perception predictions averaged for all molecules in a mixture. From the forest built in the first step, they extract the tree-embeddings which are binary vector representations containing one element per node of a tree for each tree in the forest. A value of 1 indicates that the sample traverses the corresponding node when tracing its path down the tree. If desired, one can also employ weights based on node size and dimensionality reduction techniques to further process the binary representations.

**Baseline: first leaderboard submission.** For their first submission to the leaderboard, the approach was kept relatively simple. Only the provided molecule features were used (dragon, mordred, and morgan). Each mixture was represented by averaging the feature vectors of each of its molecules (hence making the invalid assumption that molecules mix additively). A mixture pair was represented by the absolute differences in feature values for the two mixtures. Then, a random forest was trained (no tuning) on these mixture pair feature vectors, with the distance as an outcome.

**Final submission.** The final submission was based on the molecule perception embeddings described in the previous section combined with diverse representations for mixtures and mixture pairs.

**Final molecule representation.** Each molecule was represented by the 128 values of the perception embeddings concatenated with 30 principal components determined from all molecules in the train, leaderboard and test sets.

**Mixture representation.** Consider a mixture of n molecules where each molecule has m features. Each molecular feature thus presents a distribution of n values for the given mixture. They then calculate s statistics for each molecular feature, so that the mixture is represented by a total of m $\times$s numeric attributes. The statistics employed were: the arithmetic mean, standard deviation, minimum, maximum, harmonic mean, geometric mean, square mean, skewness, kurtosis, the Shannon's entropy, interquartile range, and 10%, 25%, 50%, 75% and 90% percentiles. The procedure results in a total of 2.686 mixture features.

**Mixture pair representation.** For a pair of mixtures, they consider each mixture feature in isolation. Between each pair of mixture descriptors, they calculate the mean, product, absolute difference, minimum and maximum values. Furthermore, for each molecule feature, they obtain the Mann-Whitney U statistic comparing the two mixtures. Finally, they also add a binary feature indicating whether the mixture pair originates from the Bushdid dataset, since this dataset was the only one with targets derived from the triangle discrimination test. In total, each mixture pair is represented by 13,589 features.

**Final estimator.** The final predictions were generated by an ensemble of Extremely

Randomized Trees. 1000 trees were used, and the remaining hyperparameters were set to the default values of Scikit-Learn 1.5.1. Random Forests and Gradient Boosting Machines were also investigated, with slightly worse results.

##### 1.3.2 Summary

In summary, belfaction team explored a variety of methods for representing molecules and mixtures of molecules in machine learning tasks, with the final goal of predicting perceived olfactory similarity between mixtures. They observed similar results for different molecule embedding techniques, and the most important factor to improve predictive performance was ensuring a descriptive representation of mixture pairs. Their best results were obtained by extracting a diverse set of statistical descriptors from the distributions of molecular features in each mixture. Moreover, several tree-ensemble techniques were investigated as the final predictor, with Extremely Randomized Trees resulting in the best predictive performance. The need for a detailed description of mixture pairs reflects the complex nature of molecular interactions that result in olfactory perception. Further ways of representing such interactions thus constitute interesting paths for future research. Furthermore, their methods assumed a transductive setting, in which all mixtures and molecules are known in the training step and only new mixture interactions are unknown. The performance of these techniques for completely new mixtures and compounds is another topic of future investigation. The source code of belfaction model is available at: https://gitlab.kuleuven.be/u0131222/dream2024.

#### 1.4 UMich CASI

Motivated by the use of semantic descriptors from the Keller dataset [5, 21] to determine odor mixture discriminability [6], UMich CASI team hypothesize that utilizing appropriate intermediate information can enhance the model's performance compared to directly employing chemoinformatic features. Building on this baseline method, they explored several improvements for each step.
In Step 1, they recognized that the Keller model [5], trained on 338 molecules, might have limitations. Therefore, they used open-POM [9, 10] which is trained on a larger dataset of single molecule odorants. For Step 2, they hypothesized that taking a weighted average of single molecule embeddings might be more effective than directly taking the average or maximum, leading them to employ exponential transformation. In Step 3, they proposed that ensemble learning could better fit the model, and thus experimented with three boosting methods and random forest.

##### 1.4.1 Methods

As illustrated in the pipeline illustration in Figure S3, the primary methodologies employed are graph neural networks and boosting. Specifically, they used a message passing neural network and CatBoost. The team leveraged models pre-trained on a larger dataset [9, 22] and weighted the molecules in the intersection of pre-trained data and competition data more heavily. Recognizing the significance of intensity for

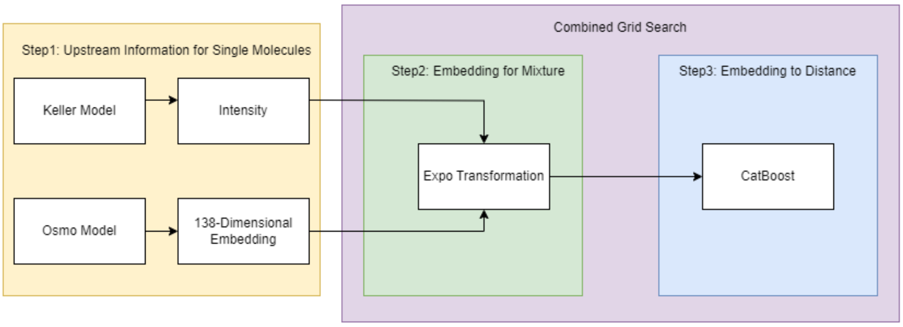


**Fig. S3** Overall pipeline of UMich CASI team approach. The three steps respectively involve obtaining pre-trained embeddings for single molecules as upstream information, using exponential transformation to derive the embedding for mixtures, and finally employing a boosting method to learn the perceptual distance

single molecules and its potential importance in mixtures, they used LassoCV to predict intensity.

**Data pre-processing.** Initially, they observed a potential distribution shift between the training set and the leaderboard set, with differences in valid Pearson coefficient performance reaching up to 0.2. Following recommendations from a discussion thread and a report [7], they utilized a new training set. Although the overall performance did not drastically improve, the data distribution shift was mitigated, reducing the difference in valid Pearson coefficient to within 0.05. This change provided more reliable parameter combinations for the final submission.

**Embeddings for single molecules.** They employed two approaches to obtain embeddings based on semantic descriptors and intensity: a message passing neural network (MPNN) and LassoCV. For the MPNN, they identified 162 molecules in the new training, leaderboard, and test sets. There were 139 molecules in the intersection of these 162 molecules and the 4983 molecules in the curated GS-LF dataset. They weighted these 139 molecules by a factor of two during training to facilitate better transfer of the pre-trained model. They used 5-fold cross-validation without a separate test set, as the MPNN was only for obtaining single molecule embeddings in Step 1. This pre-trained model provided 138-dimensional embeddings for all 162 molecules, which served as input for the next step. For LassoCV, they applied 10-fold cross-validation to predict intensity.

**From embeddings of single molecules to embedding of mixtures.** They used exponential transformation with a parameter $\tau$. A larger $\tau$ resulted in an average-like embedding, while a lower $\tau$ emphasized the molecule with the highest predicted intensity in the mixture. This approach is validated in previous studies [4]. The transformation is detailed below:

Exponentiation with temperature scaling:

$$\text{exp\_numbers}[i] = e^{\frac{\text{numbers}[i]}{\tau}}$$

Normalization:

$$\text{weights}[i] = \frac{\text{exp\_numbers}[i]}{\sum_j \text{exp\_numbers}[j]}$$

Complete formula:

$$\text{weights}[i] = \frac{e^{\frac{\text{numbers}[i]}{\tau}}}{\sum_j e^{\frac{\text{numbers}[j]}{\tau}}}$$

**Embedding to distance.** They experimented with CatBoost, XGBoost, LightGBM, and RandomForest, using the same $n_{\text{estimator}}$ settings and a combined grid search with $\tau$. The $n_{\text{estimator}}$ ranged from 200 to 500, and $\tau$ ranged from 1 to 40, with 10-fold cross-validation. They also tried several engineering methods for each 138-138 embedding pairs, exploring p-th difference (from 1 to 5) and direct concatenation. CatBoost proved to be the best model based on both the leaderboard set and the validation part of the training set.

**Data augmentation.** They tested two data augmentation approaches, both of which use a user determined threshold.

1. ***Random Approach:*** For each mixture pair, we selected a molecule and substituted it with another molecule with the highest cosine similarity (above the threshold) based on the weighted GNN embedding. If no molecule met the threshold, the pair was excluded.
2. ***Iterative Approach:*** Similar to the random approach, but iteratively changed the selected molecule until a suitable substitute was found or all options were exhausted.

where A' and B' represent the augmented mixtures of A and B. Experiments were conducted on both (A, B), (A', B) and (A, B), (A', B), (A, B'), (A', B'). Thresholds were set as follows:

- Random $\{(A, B), (A', B)\}$: 0.92
- Random $\{(A, B), (A', B), (A, B'), (A', B')\}$: 0.97
- Iterative $\{(A, B), (A', B)\}$: 0.99
- Iterative $\{(A, B), (A', B), (A, B'), (A', B')\}$: 0.997

The new training set size ranged from 800-900, with 300-400 augmented pairs.

##### 1.4.2 Summary

In summary, UMich CASI team gained the insight that increasing the weight of intersected molecules during the pre-training phase generally improves performance, outperforming the original ensemble embedding approach. In addition, they observed that while lower $\tau$ values (below 10) yielded the best performance on the training set when specific augmented datasets was used, their performance dropped significantly on the leaderboard set, indicating that this situation is not a common occurrence. Models with $\tau$ values above 30 performed more consistently, indicating that using the average is a robust estimation method. Furthermore, before updating the training set, data augmentation methods enhanced model robustness. However, as the distribution shift was mitigated, the benefits of data augmentation diminished, leading to the abandonment of this approach. The source code of UMich CASI model is available at: https://github.com/YikunHan42/DREAM-Olfactory-Mixtures-Prediction-Challenge-CASI.

### 1.5 PL21

Understanding the discriminability of odor mixtures is a complex challenge that integrates various domains such as chemistry, biology, and sensory perception. By leveraging molecular descriptors and perceptual data, PL21 team aim to predict how similar or dissimilar different odor mixtures will be perceived.

###### 1.5.1 Methods

They employed Lasso regression to model the discriminability of odor mixtures. The model inputs included the squared differences of physicochemical properties and perceptual features of the mixtures. For physicochemical properties, they utilized normalized log-transformed Dragon descriptors averaged across compounds of the mixture. The perceptual features followed the approach established by Dhurandhar et al.[6]. They found that the fraction of shared compounds between mixtures is a major determinant of discriminability. However, the target mixtures in the test set do not share compounds. Therefore, compound (dis)similarity was integrated by restricting Lasso regression training to mixtures that did not share any compounds from the dataset by Bushdid et al. [8]. All input variables were Box-Cox transformed to stabilize variance and make the data more suitable for linear modeling. The output standard deviation was reduced by a factor two while keeping the mean.

###### 1.5.2 Summary

In summary, PL21 team integrated molecular descriptors and perceptual features through Lasso regression. The inclusion of compound similarity considerations, particularly for mixtures without shared compounds, allows their model to remove a major discriminability signal that is not present in the test set.The source code of PL21 model is available at: https://github.com/Satarifard/DREAM-olfactory-mixtures-prediction-challenge/tree/main/Models/PL21.

##### 1.6 Tamarin

Modeling studies have shown that intuitive ways to compare molecular representations, such as the cosine similarity of combined features, can readily serve as good predictors for olfactory perceptual distance data (Pearson correlation of $\sim$0.42). Based on that observation, Tamarin team hypothesized that a systematic selection of molecular descriptors combined with conventional models from data science has the potential to further increase these scores. To account for the limited dataset size in this competition, they have opted for two design choices. For the molecular descriptors, they implemented a feature selection algorithm to reduce the size of the mixture representation. For the model class, they focused on compositions of decision trees [23] that have the advantage of performing better on smaller datasets such as those compiled for the challenge. They show that their approach outperforms previous models based on a single representation of molecules, embracing olfaction as a complex process involving physical, chemical, and perceptual representations of individual odorant molecules that are jointly perceived as mixtures.

##### 1.6.1 Methods

The team modeled perceptual distances between odorant mixtures based on the properties of their constituent molecules. For each molecule, they generated features based on the information that may be relevant to their perceptual representation including semantic description, structural features, and physicochemical properties. They combined molecule features to form joint features for mixtures and used them to train compositions of decision trees to predict the perceptual distances between these mixtures.

**Data proofreading.** They used the four datasets (Snitz-1, Snitz-2, Ravia-4, and Bushdid [3, 4, 8]) provided with the challenge. To account for the identified inconsistencies in the data, they performed the data proofreading and correction as follows. They started by finding invalid CIDs that they identified as erroneously concatenated pairs of CIDs. They then checked for repeated CIDs within individual mixtures and found an erroneous mixture likely formed as a result of the concatenation of a pair of mixtures due to a missing closing bracket in the original dataset (Ravia's [4]). Finally, they have identified 13 mismatched CIDs in the processed version of another dataset (Bushdid's [8]) and corrected the identified errors by matching the challenge mixtures with the original data.

**Physicochemical features for molecules.** To use the challenge molecules' physicochemical properties, they considered both molecular descriptor data (Mordred [24, 25]) and molecular fingerprint data (Morgan [26], Atom Pairs [27], and Topological Torsions [28]). As there are two versions of Mordred software [24, 25] that produce different outputs, they tested both versions of the resulting features. They reduced the dimensionality of the molecular descriptors with the quadratic programming algorithm using a weighted MSE distance between mixtures. As the molecular fingerprints' dimensionality can be arbitrarily defined, they also tested the fingerprints of different dimensionality (50, 100, 200, and 4096) and quality (binary versus frequency counting).

**Semantic features for molecules.** To account for the odorants' perceptual properties, they used the Leffingwell dataset [22] and DeepNose [29] features. DeepNose features, the additive 96-dimensional vectors encoding structural and perceptual properties of individual odorant molecules, were obtained by training a convolutional neural network model on raster images of molecules to predict their perceptual properties using the Leffingwell dataset as described before [29]. They evaluated the DeepNose features for all competition-related CIDs. They separately retrieved the Leffingwell descriptors for the same set of all challenge-related CIDs, appending them with DeepNose predictions wherever the ground truth data was not available (appended 18 out of 162 CIDs).

**Composite features for mixtures.** To form the inputs for our models, they designed mixture representations that combined single molecule features. To this end, they tested multiple methods of weighing and scaling: maximum, average, sum, and softmax over the individual molecular features. To represent a pair of odorant mixtures, they concatenated the feature vectors representing the two mixtures in the pair. Additionally, they appended the concatenated feature vector with engineered features that included the Euclidean distance, cosine similarity, and the angle between the feature vectors representing the two mixtures in the pair. Additionally, they used as inputs the total number of molecules in each mixture and both mixtures; they also used the number of shared molecules between the two mixtures in the pair. They included one-hot encoding for the data source for the training data and used KNN imputation for the leaderboard and test data. They augmented the input data by swapping the positions of the two mixtures in each data point while ensuring the data separation between training and testing folds.

**Model optimization.** They used the mixture pair features defined above as inputs to train random forest (RF) and XGBoost (XGB) models to predict the perceptual distances between pairs of mixtures. To instantiate the mixture features accounting for the perceptual and physicochemical properties of the molecules, they used either dense features and their combinations (DeepNose, Mordred) or sparse features and their combinations (Leffingwell, Morgan, Atom Pairs, Topological Torsions). Prediction is averaged over the augmented two copies for each pair of odor mixtures. For each model on each instantiation of the input features, they performed 3 rounds of the random search for the optimal sets of hyperparameters, then obtained the mean values and the standard deviations for the Pearson correlation and RMSE with 5 random seeds for each of the 3 optimized hyperparameter sets, and finally recorded the parameter set with the highest mean to represent the given model on the given instantiation of the features.

**Metamodels**. To combine the predictive power of their models based on dense and sparse features, they trained metamodels as follows. For combinations of (base) models trained on dense and sparse features, they used their predictions as inputs to train a metamodel (lasso/ridge linear/polynomial regression, RF, XGB, KNN) to predict the perceptual distances (averaged over augmented pairs) between mixtures of odorants. They evaluated the resulting metamodels in the same way as the evaluation on the base models.

**Model selection.** To select the model for the final submission, they performed an internal comparison of models. For each model (including metamodels), they used tenfold crossvalidation on the training dataset to compute the Pearson correlation and RMSE. They then ranked their models based on these parameters using the procedure described in the challenge. Specifically, they sampled 10% of held-out data 10000 times and, for each sample, calculated the performance of each model and the rankings of the model for both Pearson correlation and RMSE. They then averaged the ranks from both metrics to decide on the final ranks. As the highest-scoring models were represented by RF and XGB, they chose the RF model for the final submission due to the lesser propensity for overfitting the data. For the Leaderboard submission, they trained the best-ranking RF model on the entire Training set; for the Testing submission, they trained it on the combination of the Training and Leaderboard sets, which are publicly available through the combination of datasets [3, 4, 8].

##### 1.6.2 Summary

In summary, Tamarin team trained compositions of decision trees to predict the perceptual distances between odorant mixtures. Tree-based models, known for their high performance on small datasets with real-valued prediction targets and successful in the previous iteration of the DREAM challenge, have once again proven to be an adequate tool for modeling the perceptual distances between mixtures of odorants in this challenge. For molecules in the challenge mixtures, they have compiled and combined different representations including DeepNose, Mordred, Morgan, and Leffingwell features. They refined these representations with a feature selection algorithm, which has played a noticeable role in improving the performance of the models. Among different fingerprints that they tested, Morgan and Atom Pairs have led to a higher performance than Topological Torsion which, in turn, has led to a higher performance than RDKit. They have also found that frequency-based features were more beneficial for the model compared to the binary features. With the Mordred features, feature selection has proven to produce better results compared to the linear projection of high-dimensional features to low-dimensional space. Counterintuitively, perceptual data (Leffingwell), while being helpful, wasn't drastically more informative for perceptual distances than physicochemical data (Mordred and Morgan). They found that training separate models for sparse and dense features (both physicochemical and perceptual) and then combining their predictions within metamodels has led to the best predictions for the perceptual distance data. At the same time, the use of metamodels, while doubling the input information, did not improve the predictions for the perceptual distances by a large margin. While the choice of RF and XGB models here was dictated by the limited size of the perceptual distance datasets, other models have been developed for olfactory tasks featuring larger amounts of data. For example, graph and convolutional neural network models have shown their efficiency in olfactory percept prediction tasks where a few thousand data points are available. As more data emerges on the perceptual distances between olfactory mixtures, these alternative approaches may become relevant, shifting the focus of future research in the field toward large models and enabling the use of established data science approaches. The source code of Tamarin model is available at: https://github.com/Satarifard/DREAM-olfactory-mixtures-prediction-challenge/tree/main/Models/Tamarin.

#### 1.7 Excluded Models

All other submitted models were excluded either because of underperformance compared to random shuffled baseline, or they lacked open-source code for reproducibility. For the full list of models and performance distributions, see Figure S4

#### 1.8 Post Challenge Model

We built a post-challenge model focused on most important SHAP value of top-performing models drawn from the feature: open-POM logits, odor-pair graph embeddings, Dragon and Mordred descriptors, and DeepNose features. The aim was to match the Ensemble model performance with a single model. After tuning, the model matched top performing individual-team baselines that comprise the Ensemble model, and outperformed SOTA baselines (see Figure S12). Results suggest diminishing returns from further architectural and feature engineering tweaks, the primary constraint seems to be training data volume.

**Feature Selection:** We restrict inputs to the five families of features and convert

each mixture's component-level vectors into a single mixture vector by averaging over constituents. At pair-construction time (mixture A vs mixture B), we do not pass raw per-mixture vectors. Instead, for each base feature $f$ we derive three symmetric channels to ensure order-invariance and capture alignment/magnitude: $|\Delta| = |f_A - f_B|$, $\Delta^2 = (f_A - f_B)^2$, and dot $= f_A \cdot f_B$ Columns with missing feature entries are removed and remaining feature vector are minimally imputed.
**Model Training and Prediction:** We train an XGBoost regressor to minimize RMSE. Hyperparameters are optimized via an Optuna with fixed ShuffleSplit CV and consistent seeds. At inference we rebuild the same feature vectors, apply the fitted scaler, and obtain output as a single distance prediction per pair.

### 2 Olfactory Descriptor Families

The following olfactory descriptor family classification is used in Figure 3 for 138 semantic descirptors of POM model analysis.

- **Fruity**: apple, apricot, banana, bergamot, berry, black currant, cherry, citrus, fruit skin, fruity, grape, grapefruit, lemon, melon, orange, peach, pear, pineapple, plum, raspberry, ripe, strawberry, tropical, winey.
- **Floral**: floral, geranium, hawthorn, hyacinth, jasmin, lavender, lily, muguet, orangeflower, orris, rose, violet.
- **Green & Herbal**: alliaceous, aromatic, cabbage, camphoreous, celery, chamomile, cucumber, fresh, garlic, grassy, green, hay, herbal, leafy, mint, onion, radish, savory, tea, terpenic, tomato, vegetable, weedy, gassy.
- **Spicy & Warm**: anisic, cinnamon, clove, pungent, sharp, spicy, warm.
- **Sweet & Gourmand**: almond, caramellic, chocolate, cocoa, coumarinic, creamy, honey, lactonic, sweet, vanilla.
- **Woody & Resinous**: amber, balsamic, cedar, cortex, pine, sandalwood, tobacco, vetiver, woody.
- **Animalic & Meaty**: animal, beefy, cheesy, fishy, leathery, meaty, musk, sulfurous, sweaty.
- **Earthy & Musty**: earthy, fermented, mushroom, musty, potato.
- **Nutty, Fatty & Roasted**: burnt, buttery, coconut, coffee, cooked, dairy, fatty, hazelnut, malty, milky, nutty, oily, popcorn, roasted, smoky.
- **Chemical**: alcoholic, aldehydic, bitter, brandy, cognac, ethereal, ketonic, medicinal, metallic, ozone, phenolic, rummy, solvent, sour, waxy.
- **Complex**: clean, cooling, dry, natural, odorless, powdery.

**Table S1** Summary of training, test, and validation datasets.

| Dataset | Measurement Technique | Type | Molecules | Mixtures | Measurements | Ref |
|---|---|---|---|---|---|---|
| Snitz 1 | Similarity Rating | Train | 86 | 49 | 147 | [3] |
| Snitz 2 | Similarity Rating | Train | 43 | 14 | 91 | [3] |
| Ravia 1 | Similarity Rating | Train | 43 | 14 | 91 | [4] |
| Ravia 2 | Similarity Rating | Train | 43 | 14 | 91 | [4] |
| Ravia 4 | Triangle Test | Train | 49 | 120 | 50 | [4] |
| Bushdid | Triangle Test | Train | 128 | 520 | 37 | [8] |
| Random | Triangle Test | Test | 87 | 32 | 16 | This work |
| Manual | Triangle Test | Test | 80 | 40 | 20 | This work |
| Angle | Triangle Test | Test | 72 | 20 | 10 | This work |
| **Training** | Various | Train | 168 | 731 | 507 | [3, 4, 8] |
| **Leaderboard** | Various | LB | 161 | 92 | 46 | [3, 4, 8] |
| **Test** | Triangle Test | Test | 76 | 92 | 46 | This work |
| **Validation** | Similarity Rating | Validation | 76 | 59 | 50 | This work |

**Table S2** Summary of the top-performing models and Post-Challenge model.

| Team | ML Technique | Features | Train-Validation Split | Optimizer | Hyperparameter Optimization | Loss Function |
|---|---|---|---|---|---|---|
| **D2Smell** | XGBoost | Dragon chemical features, olfactory semantic descriptors | 100:0 | Gradient boosting | Optuna, 10-fold CV | RMSE |
| **UMich CASI** | CatBoost (XGBoost, RandomForest) | Pre-trained message passing neural network embeddings, exponential transformation on embeddings | 90:10 | Various | Grid search, 10-fold CV | RMSE |
| **ChemSenSim Lab** | Siamese Neural Network with Postprocessing using SVR | Graph Neural Network embeddings, combinatorial code of olfaction | 90:10 | Adam | 10-fold CV | MSE, SVR loss |
| **belfaction** | Extremely Randomized Trees | Molecules: D-MPNN embeddings + 30 PC. Mixtures: diverse statistics. Mixture pairs: mean, product, average, min., max., absolute difference, Mann-Whitney U. | 100:0 | Not applicable | 100-fold CV | MSE |
| **PL21** | Lasso Regression | Physicochemical properties, perceptual features | Not specified | Not applicable | Box-Cox transformation | Not specified |
| **Tamarin** | Decision Trees, RF, XGBoost | Physicochemical and perceptual properties, composite features for mixtures | Not specified | Various | Random search, 10-fold CV | Not specified |
| **Post Challenge Model** | XGBoost | Mordred, Dragon, DeepNose, open-POM, and Pair-model | | Gradient boosting | Feature importance analysis | Not specified |

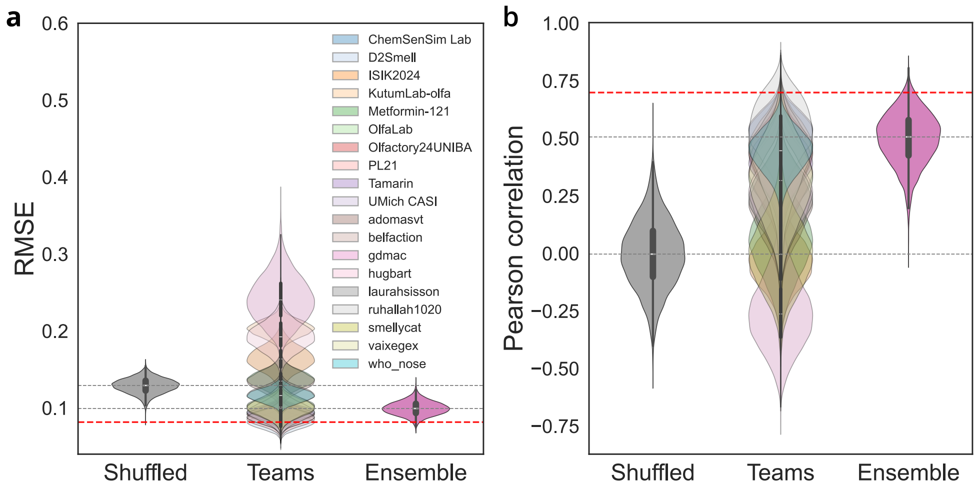


**Fig. S4 Ensemble of all submitted models. (a,b)**, Distributions of RMSE **(a)** and Pearson correlation **(b)** for all teams, the ensemble from all models, and a shuffled ground truth baseline, obtained via bootstrap sampling (n=10,000). Gray dashed lines indicate the median of the ensemble and shuffled distributions; red dashed lines denote within-subject split-half reliability (first half of the trials is used as test, and the second half as retest). Averaging these 19 models produced an ensemble model with a median RMSE of 0.1 and a Pearson correlation of 0.51.

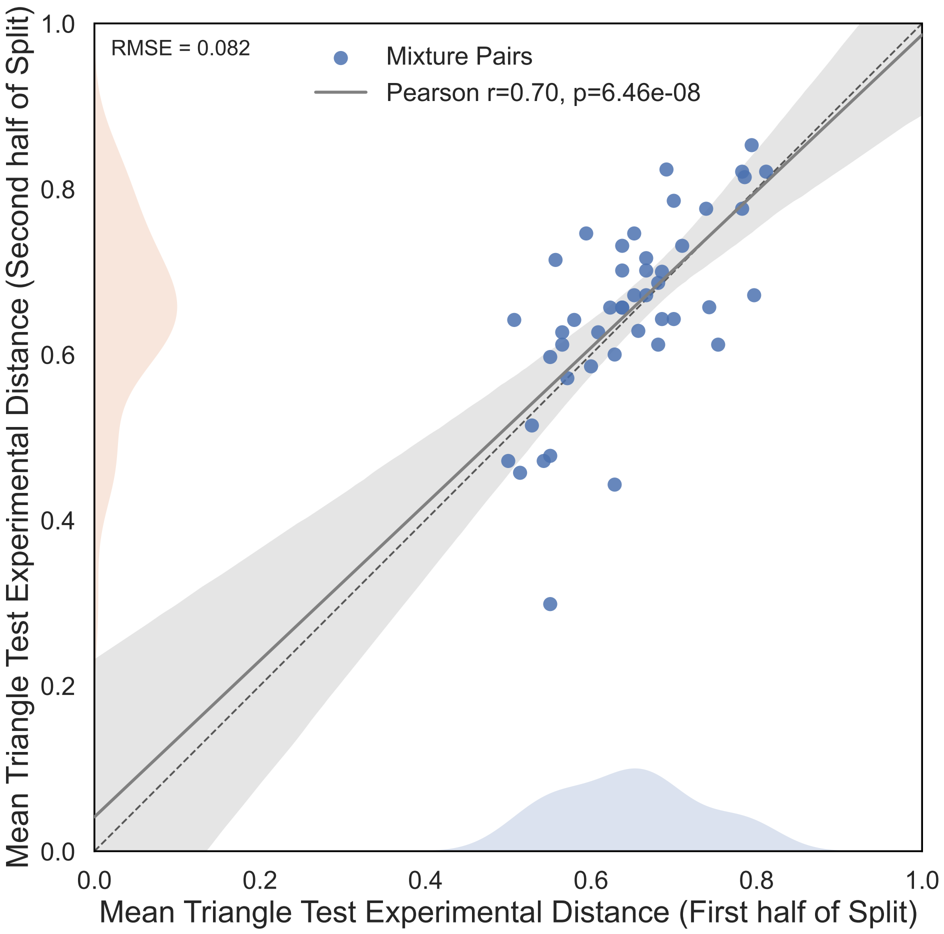


**Fig. S5 Split-half reliability of test set measurements.** The mean experimental distance values are shown for first half of split with 3188 measurements (x-axis) and second half of split with 3132 measurements (y-axis). Blue data points correspond to 46 mixture pairs in test set. Gray solid line shows the linear regression of mixture pairs with 95% confidence band. The light blue and light peach color distributions shows the distance distributions of first and second half splits, respectively.

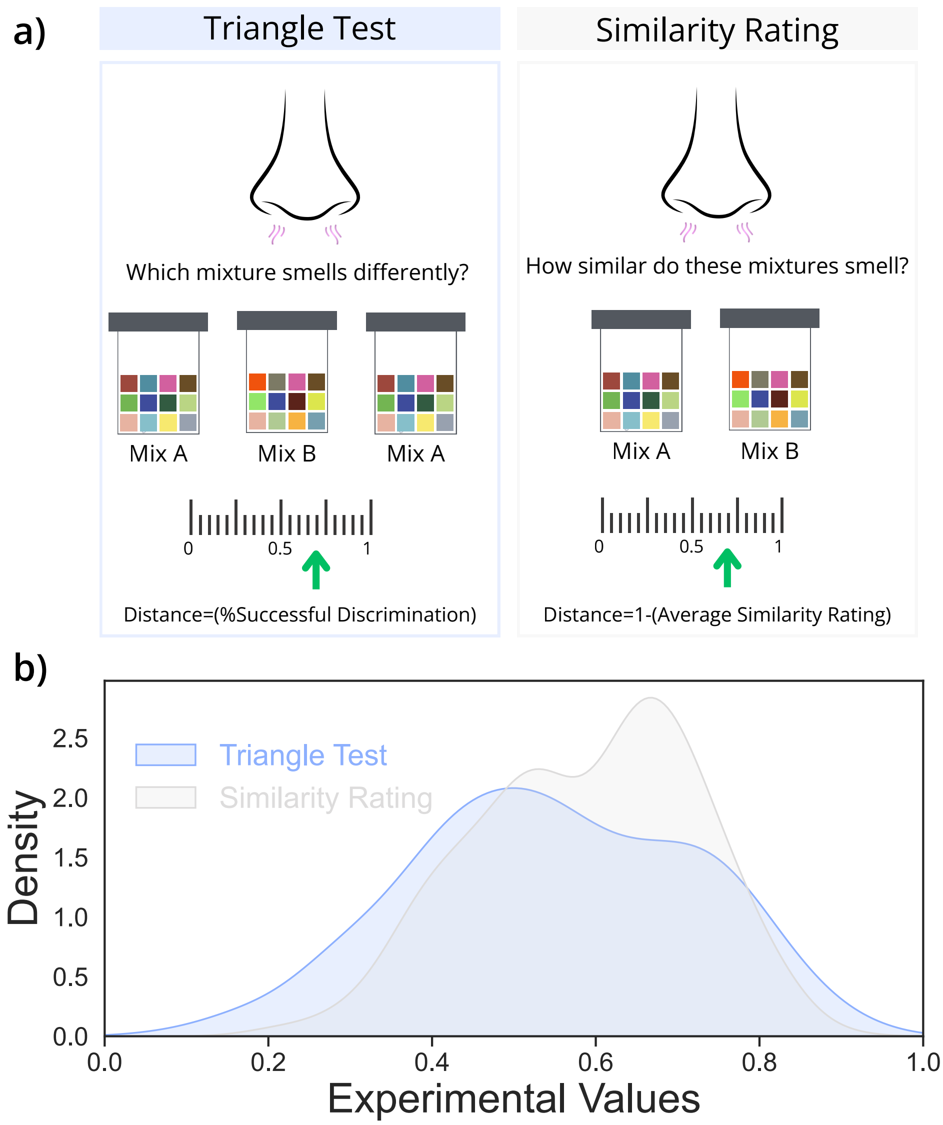


**Fig. S6 Olfactory perceptual distance measurements protocols. (a)** In triangle test, participants are tasked to select the odd odor among three vials, the olfactory mixture distance is derived from the percent correct discrimination (left). In direct similarity rating, participants are tasked to rate the similarity of two odorants, and distance is defined as 1-(average similarity rating) (right). **(b)** Training data density distribution (y axis) as a function of experimental perceptual distance (x axis) across measurement paradigms.

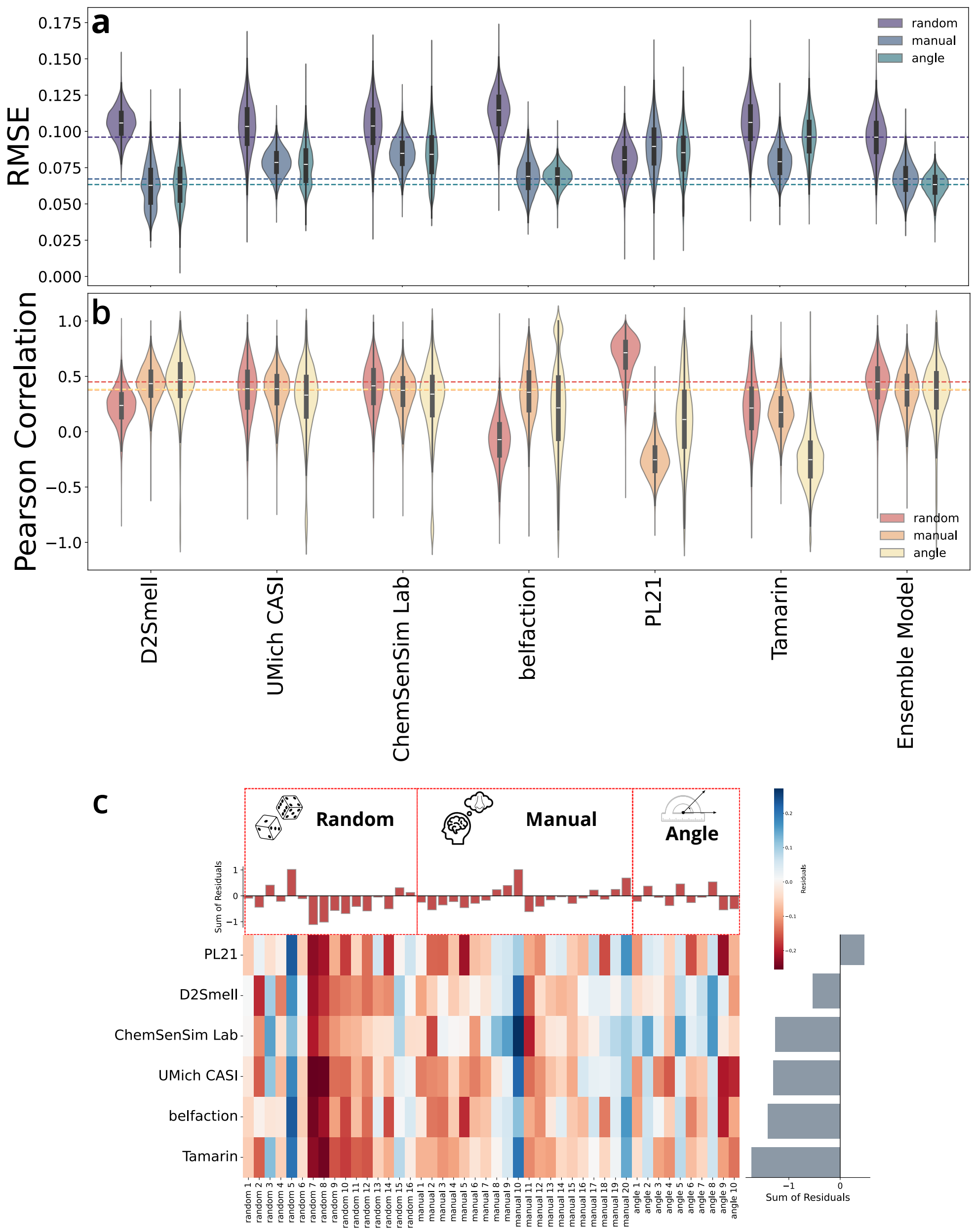


**Fig. S7 Performance across test subtypes.**

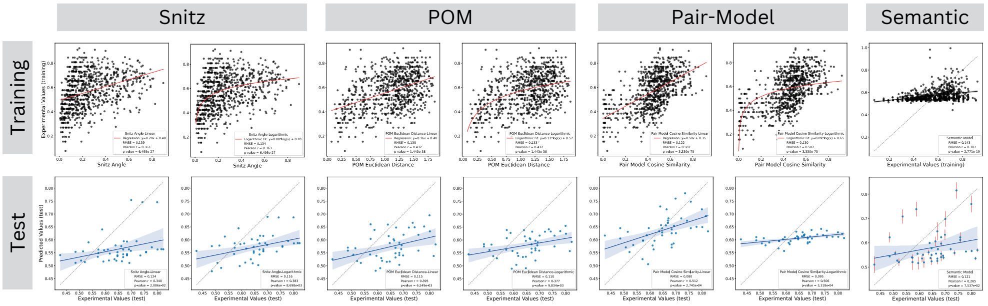


**Fig. S8 Trainings and test of state-of-the-art models.** Scatter plot show the training (top row) and test (bottom row) of SOTA benchmarks using, linear, logarithmic, or lasso regression for four models: Snitz angle [3, 4], open-POM [9, 10], an aroma chemical pair model [2], and a semantic model [6]. We fir linear and logarithmic regression to identify the best-fitting relationship (red lines in the top row) between Snitz angle, POM Euclidean distance, and Pair-model cosine similarity versus experimental distance of the training set. The best fit is then used to predict the test-set olfactory perceptual distance (bottom row). For the semantic model [6], predictions were directly generated from the open-source LASSO implementation.

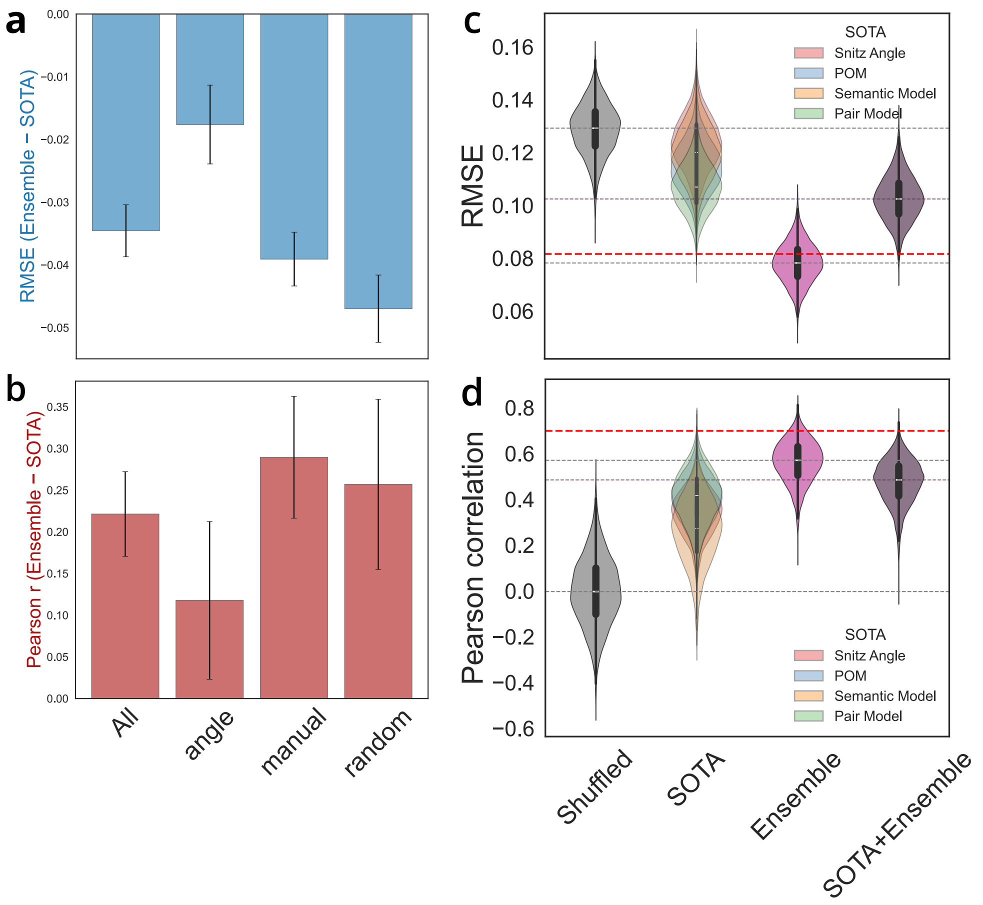


**Fig. S9 Averaging SOTA models with ensemble reduces performance**. **(a,b)** Relative change in RMSE **(a)** and Pearson correlation **(b)** when comparing the ensemble model to all SOTA models, shown across test subtypes; error bars indicate standard deviations. **(c,d)** Distribution of RMSE **(c)** and Pearson correlation **(d)** for SOTA models (linear fit), the ensemble model, and average of SOTA and ensemble model (x-axes), obtained via bootstrap sampling (n=10,000). The gray distribution represent chance-level performance from shuffled ensemble model predictions. The pink distribution represent ensemble model predictions, and violet distribution represent the average of SOTA and ensemble model predictions. All linear SOTA model are overplayed. Gray dashed lines indicate median values for the ensemble model, average of ensemble and SOTA, and shuffled baseline.

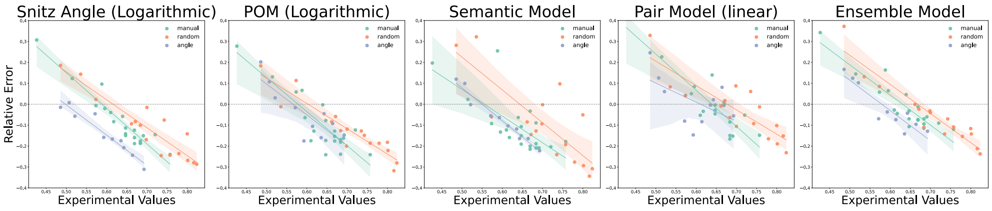


**Fig. S10 Relative error for SOTA models and ensemble model across test subtypes.** Relative error versus experimental ground-truth values for manual (green), random (orange), and angle (blue) subtypes, shown separately for four SOTA models: Snitz Angle, POM, Semantic model, pair model and the ensemble model (left to right). Solid lines (green, orange, and blue) represent linear regression fits with 95% confidence interval for each subtypes.

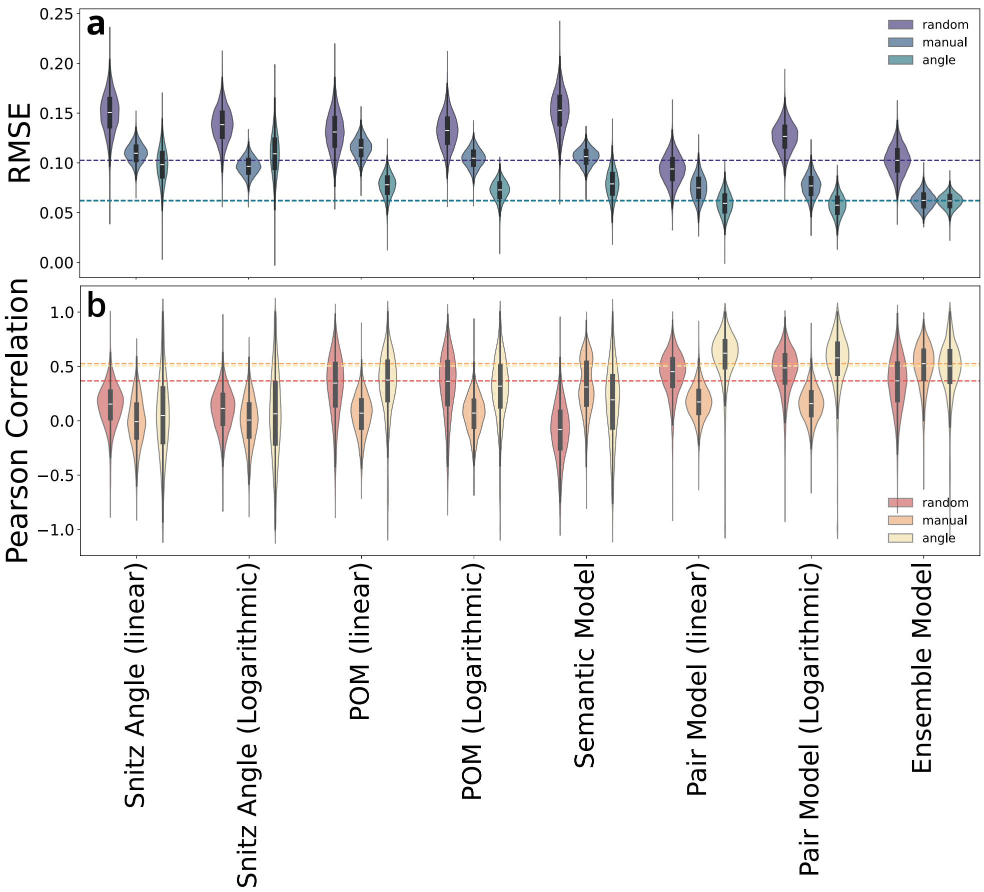


**Fig. S11 SOTA performance by test subtypes. (a,b)** Distribution of RMSE **(a)** and Pearson correlation (y axes)**(b)** for SOTA and ensemble model (x-axes), obtained via bootstrap sampling (n=10,000). Dashed lines indicate the ensemble median for each sub-types.

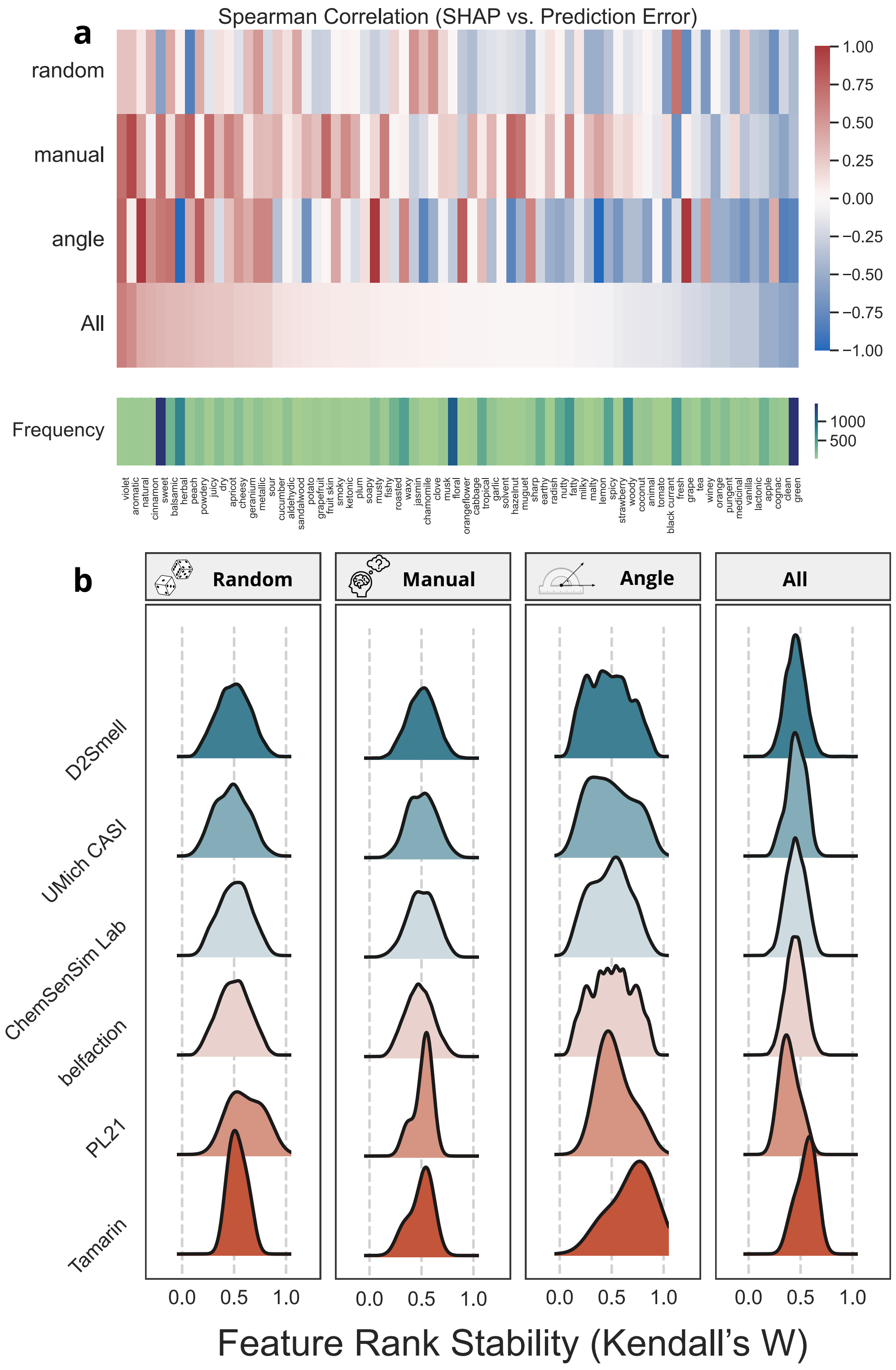




**Fig. S12** **SHAP value analysis (a)** Heatmap of Spearman's rank correlations between label SHAP values and prediction errors of POM semantic labels across test-set subtypes (top heatmap), and heatmap of label-occurrence frequency in the GoodScent and Leffingwell dataset used for training open-POM [10] (bottom heatmap). **(b)** Feature rank stability, quantified using Kendall's W, across test-set subtypes and teams.

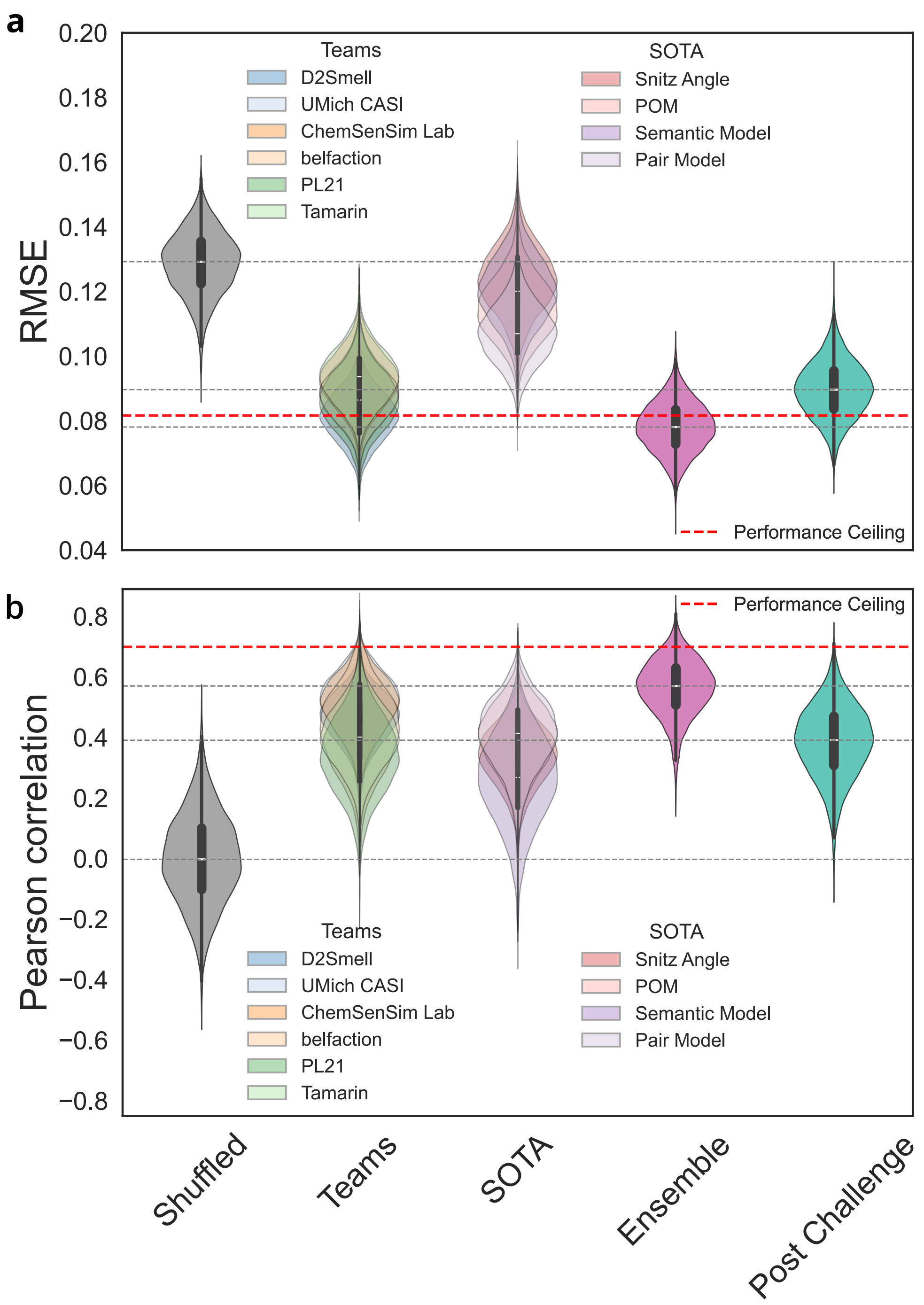


**Fig. S13 Post Challenge Model on test-set**. **(a,b)** Distributions of RMSE **(a)** and Pearson correlation (y axis)**(b)** for the ensemble model (pink), the six top-performing teams, SOTA models, the post-challenge model, and a shuffled ground-truth baseline (gray) obtained via bootstrap sampling (n=10,000). Gray dashed lines indicate median values for the ensemble model, post challenge model, and shuffled baseline; the red dashed line marks the across-subject split-half reliability (first half as test, second half as retest).

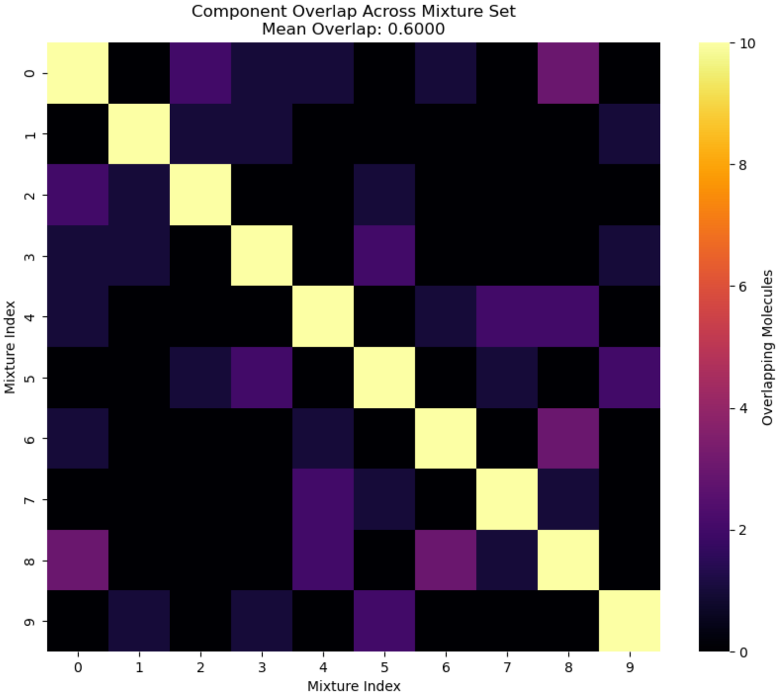


**Fig. S14 Class A Mixture Overlap**. Heatmap of molecular overlaps for 10 Class A mixtures where components overlap across mixture pairs is minimized to obtain a diverse baseline reference set.

### 3 Additional Author notes

**DREAM Olfactory Mixtures Prediction Consortium:**

Ruhallah Amandi [1], Nicola Amoroso [2,3], Loredana Bellantuono [3,4], Michele Dibattista [4], Alfonso Monaco [3,5], Ester Pantaleo [3,5], Sabina Tangaro [3,6], Gary Tom [7,8], Ella Rajaonson [7,8], Cher-Tian Ser [7,8], Sean Park [7], Stanley Lo [7,8], Brian K Lee [9], Jose M. G. Vilar [11,12], Leonor Saiz [13], Gautam Ahuja [14,15], Siddhant Poudyal [14], Bableen Kaur [14], Douluri Pushkala Devi [14], Rintu Kutum [14,15,16], Grant McConachie [17], Soroush Arabshahi [18], Saeed Karimimehr [19], Jeremy Kotlyar [19], Faraz Yazdani [20], Réka Böröcz [21], Bence Szalai [21], Adomas Malaiska [22], Victor Tarca [23], Connor Fong [23], Hugo Talibart [24], Dimitri Gilis [24], Chih-Han Huang [25], Tsai-Min Chen [26,27], Hsuan-Kai Wang [28], Jhih-Yu Chen [29], Edward S.C. Shih [30], Chih-Hsun Wu [31], Wei-Quan Fang [32], Sz-Hau Chen [33], Kuei-Lin Huang [34], Srijeet Bhattacharjee [35], Malay Bhattacharyya [35], Sergey Shuvaev [36], Cyrille Mascart [36], Khue Tran [36], and Alexei Koulakov [36].

[1] Research Division, Compressbox, Helsinki, Finland
[2] Dipartimento di Farmacia-Scienze del Farmaco, Università degli Studi di Bari Aldo Moro, 70125, Bari, Italy.
[3] Istituto Nazionale di Fisica Nucleare, Sezione di Bari, 70125, Bari, Italy.
[4] Dipartimento di Biomedicina Traslazionale e Neuroscienze (DiBraiN), Università degli Studi di Bari Aldo Moro, 70124, Bari, Italy.
[5] Dipartimento Interateneo di Fisica, Università degli Studi di Bari Aldo Moro, 70125, Bari, Italy.
[6] Dipartimento di Scienze del Suolo, della Pianta e degli Alimenti, Università degli Studi di Bari Aldo Moro, 70125, Bari, Italy.
[7] Department of Chemistry, University of Toronto, Ontario, Canada.
[8] Vector Institute for Artificial Intelligence, Toronto, Ontario, Canada.
[9] Independent.
[1]0 Department of Chemical Engineering and Applied Chemistry, University of Toronto, Ontario, Canada.
[11] Biofisika Institutua (CSIC, UPV/EHU), University of the Basque Country, P.O. Box 644, 48080 Bilbao, Spain.
[12] IKERBASQUE, Basque Foundation for Science, 48011 Bilbao, Spain.
[13] Department of Biomedical Engineering, University of California, Davis, California, USA.
[14] Koita Center for Digital Health, Ashoka University.
[15] Department of Computer Science, Ashoka University.
[16] Trivedi School of Biosciences, Ashoka University.
[17] Boston University.
[18] Columbia University.
[19] New York University.
[20] The Rockefeller University.
[21] Turbine AI.
[22] Volatile AI.

[23] Huron High School, Ann Arbor, Michigan, USA.
[24] Computational Biology and Bioinformatics, Université libre de Bruxelles, 1050 Bruxelles, Belgium.
[25] Department of Data Science, ANIWARE, Taipei, Taiwan.
[26] Graduate Program of Data Science, National Taiwan University and Academia Sinica, Taipei, Taiwan.
[27] Research Center for Information Technology Innovation, Academia Sinica, Taipei, Taiwan.
[28] Independent Researcher.
[29] Graduate Institute of Biomedical Electronics and Bioinformatics, National Taiwan University, Taipei, Taiwan.
[30] Institute of Biomedical Sciences, Academia Sinica, Taipei, Taiwan.
[31] Interdisciplinary Artificial Intelligence Center, National Chengchi University, Taipei, Taiwan.
[32] Center for Drug Evaluation, Taipei, Taiwan.
[33] Investment  Wealth Management, FCC Partners Inc., Taipei, Taiwan.
[34] Chung Shan Medical University Hospital, Taichung, Taiwan.
[35] Indian Statistical Institute, Kolkata, India.
[36] Cold Spring Harbor Laboratory, Cold Spring Harbor, NY, USA.
